# Clinico-genomic features predict distinct metastatic phenotypes in cutaneous melanoma

**DOI:** 10.1101/2025.01.24.633441

**Authors:** Tyler J. Aprati, Chi-Ping Day, Daniel Lee, Alexander Pan, Justin Jee, Giuseppe Tarantino, Michael P. Manos, Hannah Faulkner, Marta M. Holovatska, Karam Khaddour, Catherine H. Feng, Kelly P. Burke, Marc Glettig, Zoe Weaver Ohler, Rajaa El Meskini, Genevieve M. Boland, F. Stephen Hodi, Rizwan Haq, Alexander N. Shoushtari, Nikolaus Schultz, Jeffrey J. Ishizuka, Alexander Gusev, Maryclare Griffin, Kenneth L. Kehl, David Liu

## Abstract

Metastasis drives mortality and morbidity in cancer. While some patients develop broad metastatic disease across multiple organs, others exhibit organ-specific spread. To identify mechanisms underlying metastatic organotropism, we analyzed clinico-genomic data from over 7,000 patients with metastatic cutaneous melanoma in three independent cohorts (one primary discovery and two validation cohorts including a nationwide electronic health record-derived deidentified database), leveraging machine learning approaches to clinical data. We found that female sex and increased tumor mutational burden associate with decreased metastatic potential, while older age associates with increased lung and adrenal metastases. Using unsupervised analyses, patients clustered into five metastatic patterns: a “highly metastatic” cluster characterized by involvement of many organs, a “low metastatic” cluster characterized by few metastatic sites (mostly lymph node metastases), and three additional clusters each characterized by metastasis to specific sites (brain, lung, liver). Mutations in *B2M* and *PTEN* associated with increased overall metastatic potential. *PTEN* mutations were also associated with brain metastases but were enriched only in the “highly metastatic” cluster and not the brain-specific cluster. Mutations in *GNAQ* or *GNA11 (GNA)* associated with increased liver metastasis. To validate this association, we tested and demonstrated liver tropism in two GNA-mutant genetically engineered cutaneous melanoma mouse models of metastasis. Overall, our study elucidates distinct phenotypes of metastasis in patients with melanoma and identifies novel clinical and genomic associations that illuminate the drivers of clinical metastatic organotropism.

## Introduction

Metastasis is a hallmark of cancer and the primary driver of cancer mortality and morbidity in solid tumors^1^. The “seed-soil” hypothesis posits that interactions between tumor-intrinsic features and organ-specific microenvironmental factors enable metastases to form^2^. The propensity of a cancer to metastasize to specific organs, a process known as metastatic organotropism, differs by tumor histology^3^. While some cancers tend to metastasize to specific sites (e.g. prostate cancer to the bone, colorectal cancer to the liver and lung), others (e.g. melanoma, non-small cell lung cancer, clear cell renal carcinoma) display large heterogeneity between patients, with the capacity to metastasize to virtually any organ^4^. To this end, preclinical studies have unraveled mechanisms of metastasis^1,5^ and nominated mechanisms of site-specific metastasis in certain settings^6,7^, yet the clinical patterns and drivers of metastasis in patients are less well-understood, despite clear data that site-specific metastases have prognostic and predictive significance^8,9^. One challenge is that standardized ascertainment and annotation of organ-specific metastasis in patients is difficult; early studies used sites of metastasis determined from autopsy studies^10–13^, but this represents cross-sectional data at late stage of disease without accompanied molecular data to analyze associated biological mechanisms.

Recently, machine-learning methods have enabled development of clinico-genomic datasets at scale with highly accurate annotations of metastatic sites compared to prior methods^14–17^. These studies focused on site-specific analyses, examining associations of specific gene mutations with site-specific metastasis across and within tumor histologies (e.g. RB1 mutations associated with CNS and liver metastases in lung adenocarcinoma^16^). However, the analysis of heterogeneity in patterns of metastatic disease at the patient level in these cohorts has been limited. Some patients have widely metastatic disease (e.g. spread to the liver, lung, lymph nodes, brain), while others only develop metastasis in a limited number of organs (e.g. brain). We hypothesized that the biological mechanisms driving metastasis, mediated in part by genomic mutations and patient clinical features, differ between these groups of patients and drive different patterns of site-specific metastasis.

To test this hypothesis, we perform a rigorous analysis of temporal metastasis patterns and their genomic and clinical correlates in over 7000 cutaneous melanoma patients from 3 independent cohorts (one primary discovery, two as validation cohorts) with independent platforms for longitudinal metastatic site abstraction and genomic sequencing. Cutaneous melanoma has broad and heterogeneous clinical metastasis patterns, in addition to well-characterized and clinically relevant genomic drivers, making it particularly well-suited for this analysis, and has not been included in prior studies^16^ of machine-learning derived clinico-genomic cohorts.

From unsupervised analyses, five major patterns of metastasis emerge. Two subgroups of patients are defined by high- and low-metastatic potential (number of organs with metastasis), and three subgroups are dominated by specific metastatic sites (brain, lung, and liver). We validate these patterns in an independent patient population. Using time-to-event models, we further identify clinical and genomic correlates of rapid and slow site-specific and overall metastatic potential in melanoma. Overall, female sex and higher tumor mutational burden (TMB) associated with decreased metastatic potential, while older age associates with higher rates of lung and adrenal-specific metastasis. Two alterations previously demonstrated to be associated with immunotherapy resistance, loss-of-function mutations in *B2M* and *PTEN,* are identified as the strongest predictors of high metastatic potential. Consistent with prior observations, *PTEN* mutations are associated with brain metastases in our cohort but enrich only in the group of patients with high metastatic potential; it is not enriched in the group of patients with more isolated metastasis to the brain. Further, we find distinct genomic features between brain metastases in these groups. Lastly, we identify driver mutations associated with rapid acquisition of liver metastases not previously identified in cutaneous melanoma, which we validate in an independent patient cohort and preclinical mouse models. Overall, our study provides novel insights into patterns of metastasis in patients and their relationship to genomic and clinical features in cutaneous melanoma, creating a framework to elucidate the biological mechanisms driving the heterogeneity of clinical metastasis that can be applied to other tumor types.

## Results

### Clinical and genomic characteristics of discovery cohort

We identified 881 patients with available next-generation sequencing^18^ (NGS) of their melanoma tumors from 2005 to 2023 at Dana-Farber Cancer Institute (DFCI). We collected 26,117 text imaging reports from these patients, utilizing a natural language processing (NLP) model to annotate presence of cancer and presence of metastatic sites^14,15^. After filtering for patients with metastatic cutaneous melanoma and imaging scans relevant for metastatic site annotations (Methods), our final cohort consisted of 13,033 cancer relevant scans across 455 patients (Fig. 1a-b). Patients with cutaneous melanoma had a median follow up of 19 months and 22 annotated reports. The median age of patients was 64 years old at time of first radiographic evidence of cancer, and 64% of the patients were male. Annotated sites of metastasis included brain, bone, lymph node (LN), lung, liver, adrenal gland, and peritoneum (Fig. 1c). For each patient, we generated time to each site-specific metastasis, utilizing the first radiographic report with cancer detected as reference time zero.

**Figure 1.**
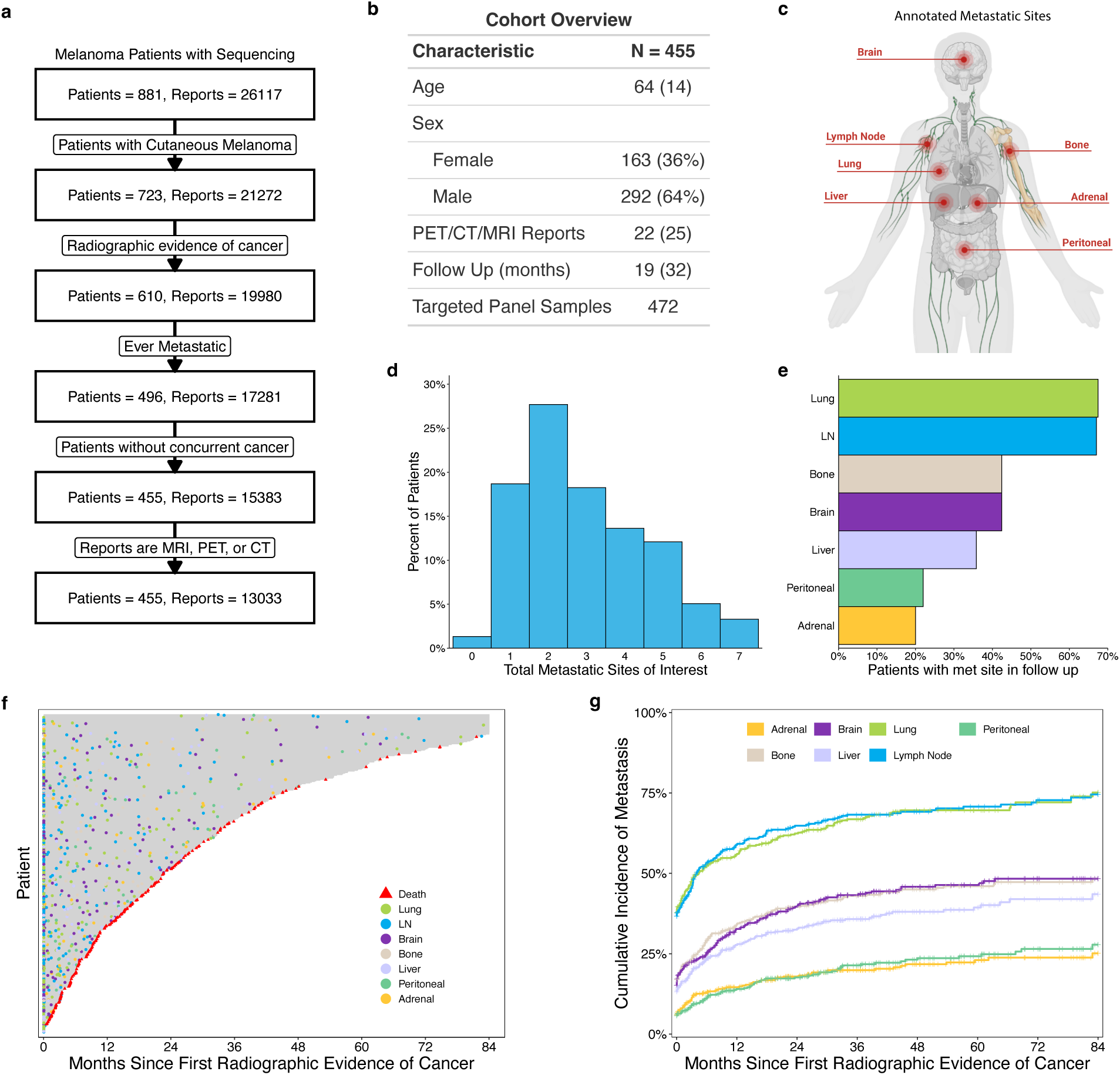
Cohort clinical characteristics and metastatic frequencies. **a.** Overview flowchart of imaging report filtering steps for cutaneous melanoma patients. **b.** Summary of cohort clinical characteristics. All statistics are presented as median (Standard deviation) or count. Follow-up time starts from date of first radiographic evidence of cancer. **c.** Schematic showing sites of metastasis annotated by the NLP model. **d.** Histogram of total distinct metastatic sites for all cutaneous patients in their lifetime. **e.** Overall frequency of metastatic sites for all cutaneous patients in the follow-up period. **f.** Event plots for cutaneous patients. Each row represents a patient, with metastatic events appearing as dots. Follow up for each patient ends with censoring or death. **g.** Cumulative incidence plot of each metastatic site for all cutaneous patients.

We next analyzed the 472 NGS panel sequenced tumors from these patients. 17 patients had more than one tumor that was sequenced, and information was aggregated for these patients (Fig. 1b) (Methods). Tumor tissue used for sequencing was primarily from metastatic (vs primary) sites and most often from skin biopsies (Supp Fig. 1a-b). Mutations in 158 genes common to all versions of the targeted panel sequencing were evaluated and further filtered so that only alterations annotated as functionally relevant were considered in the analysis (Methods). 41% had mutations in BRAF, 33% in NRAS, and 20% in NF1 (Supp Fig. 1c), consistent with previously reported driver mutation frequencies in cutaneous melanoma^19^. Mean TMB for the cohort was 21.2 Mt/Mb with a standard deviation of 19.5 Mt/Mb. Further cohort details can be found in Supplementary Data.

Patients had an average of 2.97 organs of metastasis (median 3, mode 2) in the follow-up period (Fig. 1d). Lung and lymph node were the most common metastatic sites (*∼* 70% lifetime), followed by bone, brain, and liver metastatic sites (35-45%), and peritoneal and adrenal metastatic sites (*∼* 20% lifetime) (Fig. 1e). The most common combinations of metastatic sites across patient lifetime were lung and LN, lung only, and brain only, which occurred in 9%, 7%, and 4% of patients, respectively (Supp Fig. 1d). Longitudinal metastasis data for the cohort is show in Fig. 1f. At 4 years after first radiographic evidence of cancer, we observed a cumulative metastatic incidence of 66% (lymph node), 60% (lung), 39% (bone), 35% (brain), 30% (liver), 19% (peritoneal), and 16% (adrenal) (Fig. 1g). The acquisition of metastatic sites was concordant with their lifetime frequency: lymph node and lung metastases were frequent and occurred earlier; bone, brain, and liver metastases were less frequent and occurred throughout patient follow-up; adrenal and peritoneal metastases were rare and occurred later in the disease progression.

### Association of site-specific metastasis on survival and other metastasis

We then evaluated how development of site-specific metastasis predicts overall survival and later metastasis, modeling site-specific metastases as time-dependent covariates and adjusting for total number of other metastatic sites (Fig. 2a).

**Figure 2.**
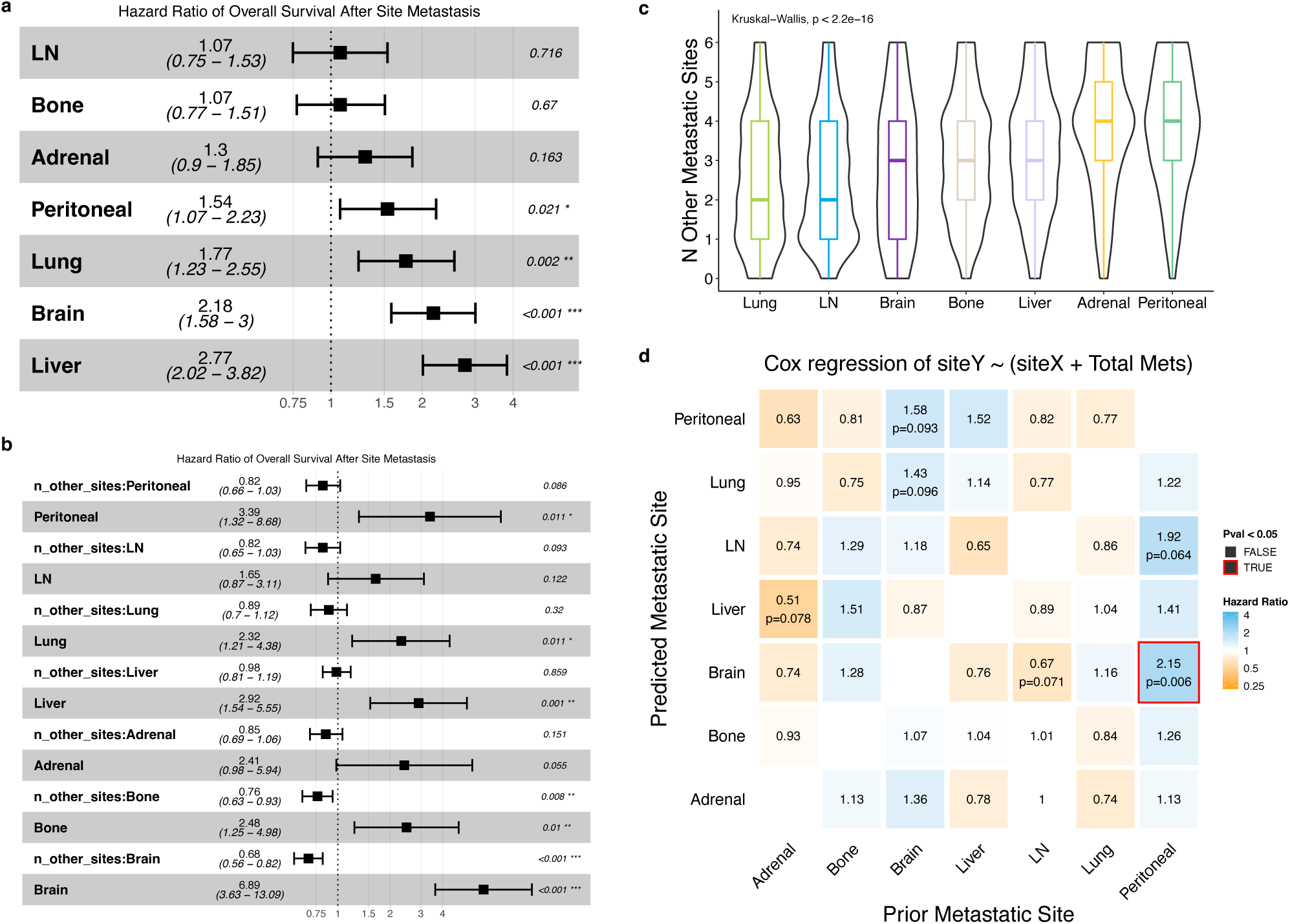
Associations of Site Metastasis on outcomes. **a.** Forest plot of multivariable cox regressions modeling the impact of each site on overall survival. Each site was run as an independent model of the form `OS site + total other sites` with a term for the site of interest and a variable for total metastatic sites. Sites were encoded as time dependent covariates, and total sites was encoded as a cumulative time-dependent covariate. **b.** Forest plot of a multivariable cox regression showing the impact of each site and site interactions with total other metastatic sites on overall survival. Each site was run as an independent model of the form `OS site * total other sites`. **c.** Violin and boxplots of total number of metastatic sites (lifetime) stratified by site. The area within each violinplot is normalized to be equivalent. **d.** Time dependent site-site associations. Squares represents the cox regression hazard ratio of the time dependent covariate prior metastatic site (X axis) with the incidence of the predicted metastatic site (Y axis). Total sites was included as a cumulative time dependent covariate in each model. Boxplots: Box limits indicate the IQR (25th to 75th percentiles), with the center line indicating the median. Whiskers show the values ranges up to 1.5 x IQR above the 75th or below the 25th percentiles. Forest plots: The center box indicates estimated HR with whiskers showing the range of the 95% confidence interval for the HR.

Development of any metastatic site was associated with worse survival, but liver and brain metastases had the worst effects on survival (HR[95% CI] = 2.77[2.02-3.82], 2.18[1.58-3.0], respectively, all p < 0.001). The associations between sites and survival and ordering of effects appear consistent with previous exploration of survival differences by first detected site of metastasis^20^.

We next tested whether the prognostic impact on survival of developing a metastatic site was dependent on the number of other metastatic sites, e.g. if the effect on overall survival differs if a metastatic site is, for instance, the first or fifth site of disease (Fig. 2b). Interestingly, liver and brain metastases were different; brain metastasis conferred a 589% increased rate of death as the first site of metastasis, while as the fifth site of metastasis only conferred a 47% increased rate of death (Brain HR = 6.89, 95% CI [3.63-13.09], p < 0.001; number of met sites:brain interaction HR = 0.68, 95% CI [0.56-0.82], p < 0.001). On the other hand, liver metastasis conferred an estimated 192% increased rate of death without statistically significant difference by the number of other metastatic sites (Liver HR = 2.92, 95% CI [1.54-5.55], p = 0.001; number of met sites:liver interaction HR = 0.98, 95% CI [0.81-1.19], p = 0.86).

We then examined the associations between site-specific metastasis and overall lifetime metastatic burden. Adrenal and peritoneal metastases were associated with the highest metastatic burden, lung and lymph node metastases were associated with the lowest overall metastatic burden, and liver, bone, and brain metastases were associated with intermediate metastatic burden (Fig. 2c, Supp Fig. 1e).

We next investigated associations between metastatic sites. We took a temporal approach, modeling the time-dependent relationship between a specific site of metastasis and subsequent development of another site of metastasis, adjusting for the total number of metastatic sites (Fig 2d, Methods). The strongest associations observed were a 115% and 92% increased rate of developing brain or lymph node metastases, respectively, in patients after development of peritoneal metastases (HR = 2.15, 95% CI [1.24-3.73], p = 0.006; HR = 1.92, 95% CI [0.96-3.81], p = 0.064; for brain or lymph node, respectively). We observed a trend that patients who had LN metastases without prior brain metastasis had a 33% lower rate of developing subsequent brain metastasis (HR = 0.67, 95% CI [0.44-1.03], p = 0.071), while the converse (brain metastasis negatively predicting subsequent development of lymph nodes) was not observed (HR = 1.18, 95% CI [0.79-1.77], p = 0.42). Other trends included 1) positive association of prior peritoneal metastasis with development of LN metastasis (HR = 1.92, p = 0.064), 2) positive association of prior brain metastases and subsequent development of lung and peritoneal metastases (HR = 1.43, 1.58, p = 0.096, 0.093, respectively), and 3) negative association of prior adrenal metastasis with subsequent liver metastasis (HR = 0.51, p = 0.078). Generally, adrenal metastasis was negatively associated with subsequent development of metastases at other sites. In contrast peritoneal metastases, a similarly low frequency site, positively associated with development of metastases at other sites. Taken together, our analysis of the effect of site metastasis shows that specific sites associate with differences in survival, total metastatic site burden, and other distant metastatic sites.

### Clinico-genomic correlates of metastatic potential

To explore overall dynamics of metastasis among all patients, we analyzed the number of organ sites with metastasis over time, utilizing the first radiographic evidence of cancer as time zero (Fig. 3a, Methods). At time zero, patients had a mean of 1.3 metastatic sites, and at 84 months, a mean of 4.3 metastatic sites. To analyze features that associated with different rates of acquiring metastatic sites, we developed a multivariable Andersen-Gill hazard model including sex, age, and TMB as covariates (Fig. 3b-c). A ten-fold increase in TMB and female sex were associated with 32% and 18% lower rates of new metastatic sites, respectively (HR = 0.68 per log_10_(TMB), 95% CI [0.58-0.8], p < 0.001; HR = 0.82 for female sex, 95% CI [0.73-0.92], p < 0.001, respectively). Age was not associated with a difference in rate of overall metastatic site acquisition. Next, we assessed the impact of functional gene mutations on the rate of new metastatic sites (Fig. 3d). In univariable analysis, patients with *B2M*, *RB1*, or *PTEN* mutated tumors exhibited a significantly higher rate of new metastatic sites, whereas patients with *EP300* mutated tumors exhibited a significantly lower rate of new metastatic sites (Fig. 3d-e, Supp Fig 3a-b).

**Figure 3.**
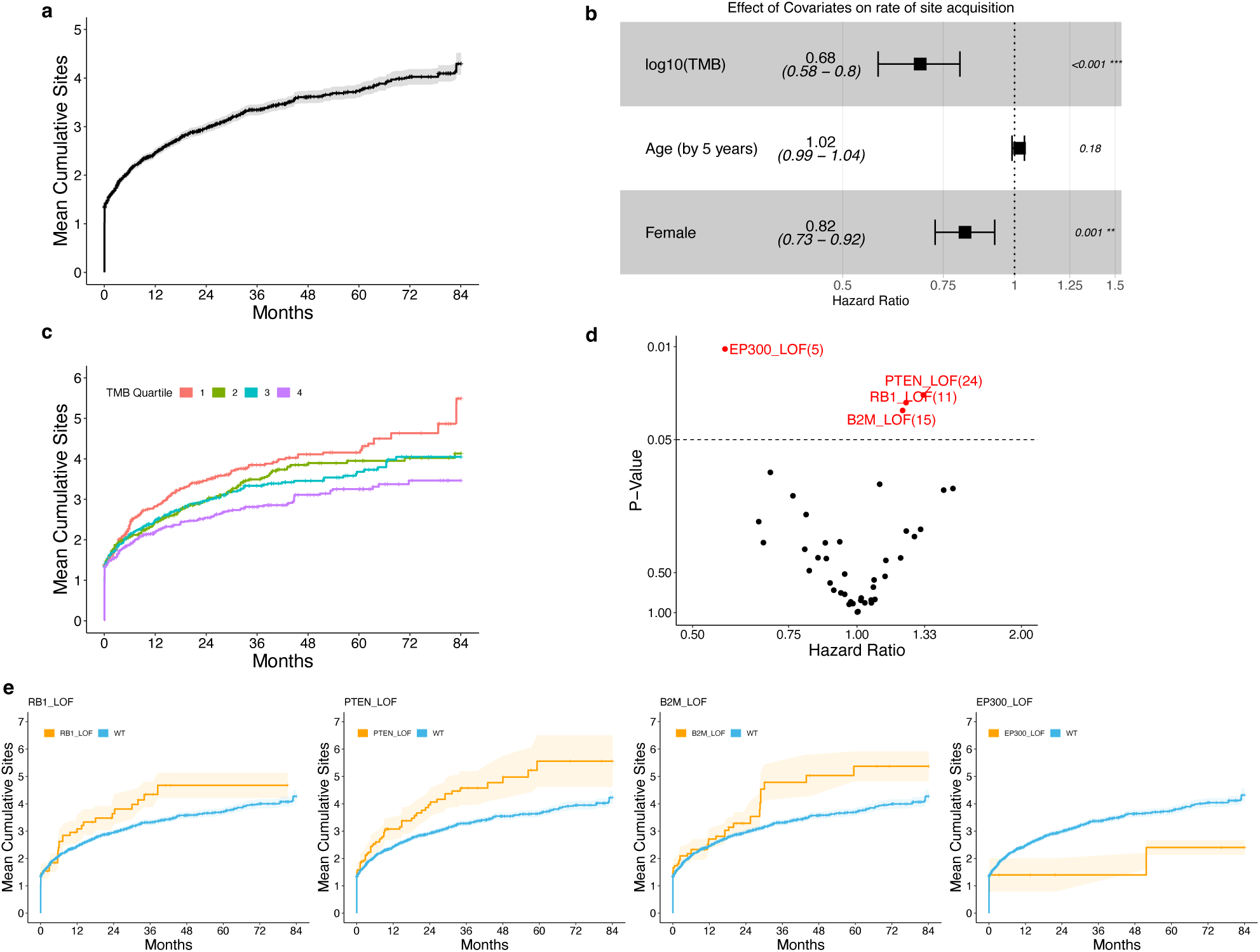
Correlates of Metastatic Potential. **a.** Estimated mean cumulative function of total distinct metastatic sites across patient follow-up. **b.** Estimated effect of clinical covariates on rate of new metastatic sites. The forest plot shows the results of a multivariable Andersen-Gill model for recurrent events. For forest plots, the center box indicates estimated HR with whiskers showing the range of the 95% confidence interval for the HR. **c.** Mean cumulative function of total distinct metastatic sites of patients stratified by TMB quartile. Quartile 1 represents the lowest TMB values. **d.** Volcano plot of univariable Andersen-Gill models for each gene. Genes where at least 5 patients were mutated are shown. **e.** Mean cumulative function of total distinct metastatic sites of patients stratified whether patients had functional mutations in select genes.

### Patterns of metastasis

Having characterized metastatic frequencies and identified features of overall metastatic potential, we then investigated patient-level patterns of site metastasis. We clustered patients by lifetime metastatic site status using a Bernoulli mixture model and selected five clusters based on AIC and median silhouette scores, observing strong evidence of clustering structure (empiric p-value of silhouette score < 0.001, Methods, Fig. 4a, Supp Fig. 4a-d).

**Figure 4.**
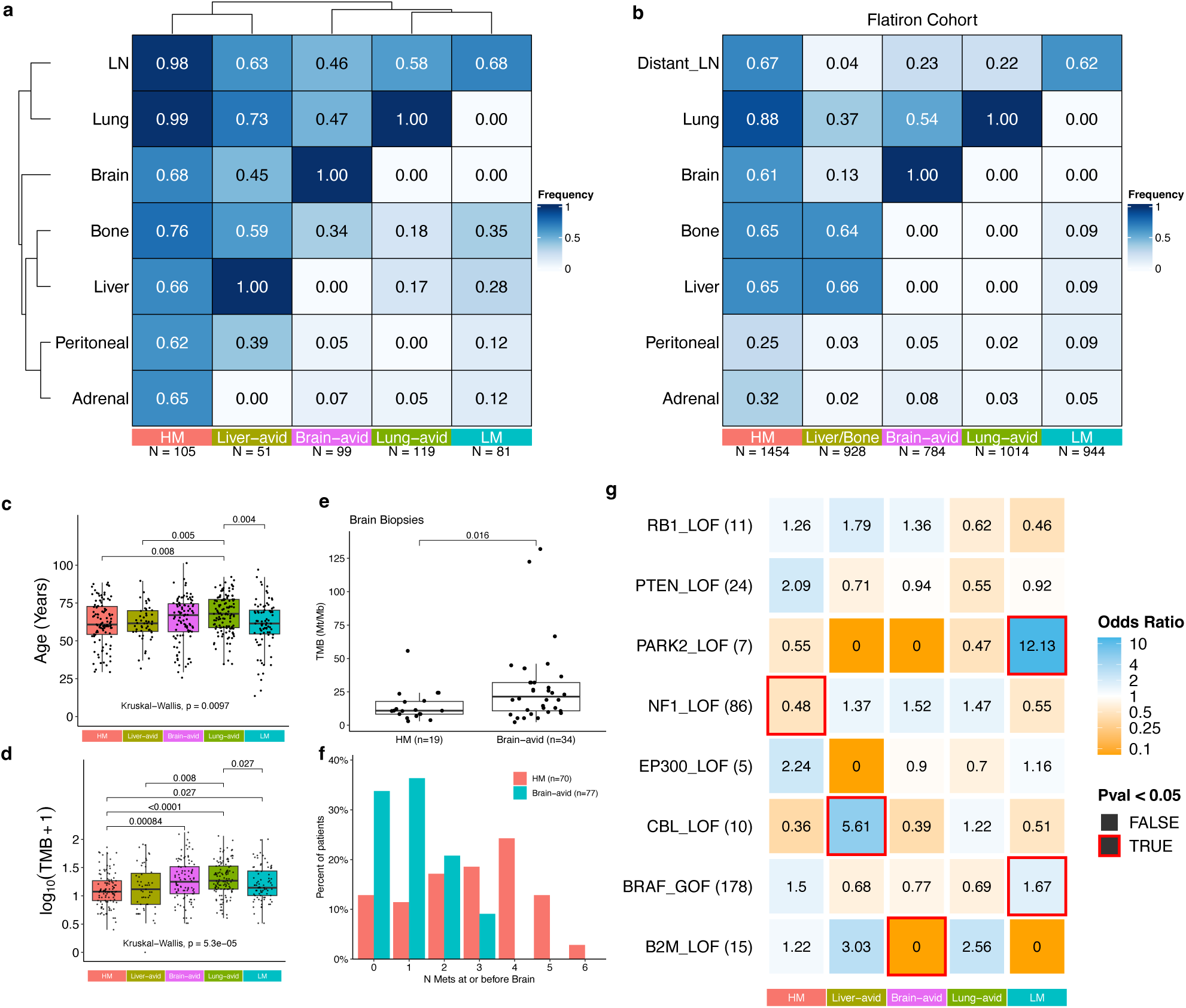
Patterns of metastasis across Cutaneous Melanoma Patients. **a.** Lifetime occurrence frequency of metastatic sites by cluster. Sites and clusters were grouped using hierarchical clustering. Numbers under each column are the total patients assigned to each cluster. **b.** Lifetime occurrence frequency of metastatic sites by cluster within the Flatiron cohort. Numbers under each column are the total patients assigned to each cluster. **c.** Boxplot of age at first radiographic evidence of cancer by cluster. **d.** Tumor mutational burden by cluster. Boxplots show log_10_TMB by cluster. **e.** Boxplot comparison for TMB values for brain biopsies from patients in the Brain-avid (n=34) or HM cluster (n=19). Test bar represents the p value of a wilcoxon rank sum test. **f.** Grouped boxplot showing the number of other metastatic sites before or at brain metastasis in patients among the Brain-avid and HM clusters. Only patient who acquire brain metastases are included. **g.** Heatmap displaying enrichment of mutations by cluster. Each square represents a Fisher test of functional mutation occurrences in the specified cluster vs. all other clusters. Red borders represent tests with P-Values < 0.05. Numbers next to gene names refer to the number of patients with that functional mutation across the cohort. Boxplots: Box limits indicate the IQR (25th to 75th percentiles), with the center line indicating the median. Whiskers show the values ranges up to 1.5 x IQR above the 75th or below the 25th percentiles.

The clusters were distinguished by metastatic burden (defined by number of organs with metastasis) and dominant sites. The “HM” cluster (highly metastatic) contained 23% of the cohort, was distinguished by high metastatic burden (mean of 5.33 metastatic sites). The “Liver-avid” (high liver) cluster contained 11% of patients and was defined by the presence of liver metastasis, along with a high frequency of bone, LN, and lung metastases (mean of 3.78 metastatic sites). The “Brain-avid” (high brain) cluster contained 22% of patients and was defined by the presence of brain metastasis and low metastatic burden (mean of 2.4 metastatic sites). The “Lung-avid” (high lung) cluster contained 26% of patients and was defined by lung metastasis and low metastatic burden (mean of 1.97 metastatic sites). Finally, the “LM” (low metastasis) cluster contained 18% of patients and was defined by the presence of lymph node metastasis and especially low metastatic burden (mean of 1.55 metastatic sites).

Interpretability and stability of these clusters was reinforced by examining how patients were distributed into clusters as the number of clusters increased using a “cluster tree.” We found that k=2 separated patients into the highly metastatic and low metastatic distinction, while at k=3 the brain-avid subset of patients emerged and remained relatively stable at all higher values of k (Supp Fig. 4e-f). At k=4, the lung-avid group appeared as a subset of the low metastatic patients, and then at k=5 the liver-avid cluster mainly came from a subset of the highly metastatic patients. Overall, the major clusters we describe remained stable at larger numbers of clusters, and they naturally followed as offshoots of the broadly high or low metastatic groups.

Next, we sought an orthogonal dataset to validate the observed patterns. We analyzed an independent cohort of 5,124 patients with metastatic cutaneous melanoma (Flatiron cohort) whose lifetime metastasis site status was annotated using a different methodology (Methods). Notably, follow up time in this cohort was shorter than our DFCI cohort (median 12, stdev 30.8 months vs. median 18.7, stdev 31.6 months respectively) with concordantly fewer metastatic sites (mean of 2.24 vs. 2.97 metastatic sites among the shared annotated sites respectively). Clustering patients from this independent cohort using the same methodology fell into the same broad categories: highly metastatic, liver/bone-avid, brain-avid with low other sites, lung-avid, and low metastasis, with a few differences (e.g. overall a lower frequency of metastatic sites, including adrenal, peritoneal, and bone metastases) (Fig. 4b, Supp Fig. 5a-e). To test if these independent cohort clusters were concordant with the original clusters, we trained a model to classify patients into the five clusters (Methods) using the discovery cohort data, then used the model to classify patients from the Flatiron cohort into these clusters (Supp Fig. 5f, Methods). We found strong associations between the similarly labeled clusters (e.g. OR for the lung-avid cluster in the independent cohort to the lung-avid cluster in the original cohort = 432.6, p < 0.001, Supp Fig. 5g), suggesting strong similarities in clustering structure.

### Genetic and clinical features of metastatic clusters

We then compared clinical features between our metastatic clusters. Age and TMB were significantly different between clusters (Kruskal-Wallis p = 0.0097, Kruskal-Wallis p < 0.0001 respectively, Fig. 4c-d). The Lung-avid cluster contained the oldest patients. The Brain-avid and Lung-avid clusters had the highest TMB (median 16.7/17.5 mutations/MB, respectively), while the HM cluster exhibited the lowest TMB with a median of 10.9 Mt/Mb. We observed a trend that patients in the HM and Liver-avid clusters were more likely to be male compared to the other clusters (Fisher Exact *p* = 0.0502, OR = 1.54) (Supp Fig. 6a). The HM and Liver-avid clusters had more imaging reports and longer follow-up (Supp Fig. 6b-d). To ensure cluster phenotypes were not confounded by these differences in follow-up, we performed landmark analysis of total metastatic sites at 12 months and found that the results were consistent with the full follow-up analysis (Supp Fig. 6e). The HM and Liver-avid clusters had the worst overall survival across clusters, while the Lung-avid and LM clusters had the longest survival (Supp Fig. 6f-g).

While patients in different clusters developed brain metastases, their clinical and disease genomic characteristics differed. Comparing brain biopsies from patients in the HM and Brain-avid clusters (n = 34, 19, respectively), brain biopsies from patients in the Brain-avid cluster had an approximately 2-fold higher TMB than brain biopsies from patients in the HM cluster (median 28.7 vs 10.9 respectively, Wilcox p = 0.016, Fig. 4e). Furthermore, the occurrence of the brain metastasis relative to other metastatic sites was lower in the Brain-avid cluster compared to the HM cluster (median 1^st^ site vs 3^rd^ site respectively, Wilcox p < 0.001, Fig. 4f).

Next, we tested if functional gene mutations were enriched in any cluster (Methods) (Fig. 4g). Mutations in *BRAF* and *PARK2* were significantly enriched in the low metastatic cluster. Mutations in *NF1* were significantly depleted in the HM cluster. Mutations in *CBL* were significantly enriched in the Liver-avid cluster. Interestingly, mutations in *B2M*, which were associated with increased overall metastatic potential, were mutually exclusive with the Brain-avid and LM clusters (OR = 0, both, p = 0.05 and 0.085, respectively).

### Clinical and Genomic Associations of site-specific metastasis

Moving from broad metastatic potential and patterns, we investigated features associated with site-specific metastasis. To address this question, we analyzed clinico-genomic predictors of longitudinal site-specific time-to-metastasis (Methods).

Broadly, results were consistent with prior associations of female sex and higher TMB with decreased overall metastasis (Fig. 5a). The strongest effect sizes of female sex and high TMB were observed with decreased rates of liver metastasis (cause-specific hazard ratio (CSHR, Methods) of 0.62 and 0.40, 95% CI [0.44-0.87] and [0.24-0.66], p = 0.005 and < 0.001, respectively, Fig 5b). Interestingly, age was not associated with overall differential metastatic rates adjusting for sex and TMB, but it was significantly associated with increased rates of lung and adrenal metastases (7% and 9% increased rate of metastasis per 5 years of age, p=0.002 and 0.03, respectively, Fig 5b). Interestingly, we observed that in some cases, a threshold value of TMB seemed to distinguish patients with different rates of site-specific metastasis; the highest quartile of TMB separated from the other three quartiles with the lowest adrenal metastasis rate, while the lowest quartile of TMB separated from the other quartiles with the highest rate of peritoneal metastasis (Fig. 5b).

**Figure 5.**
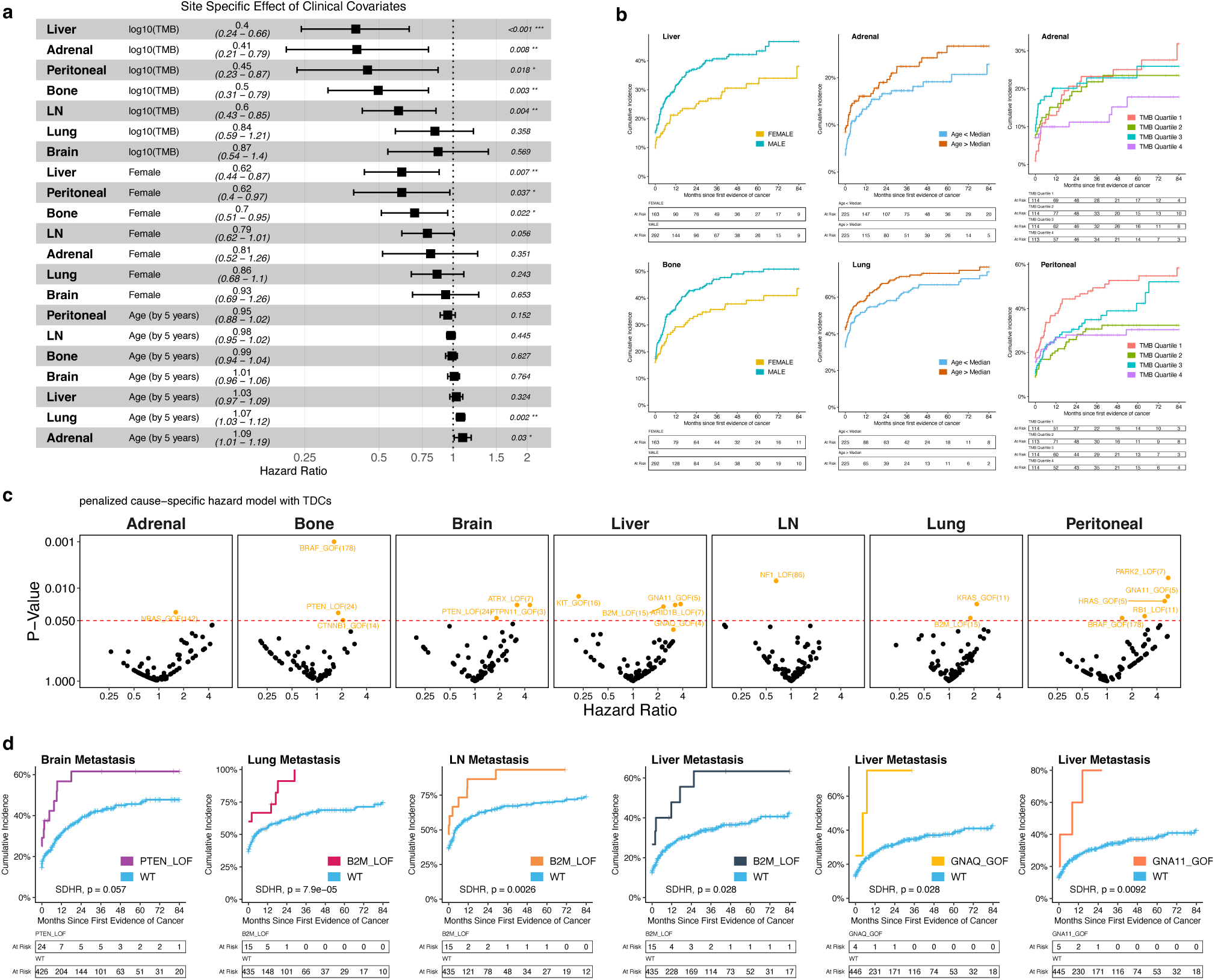
Associations of Site-Specific Metastasis. **a.** Forest plot displaying the results of multivariable cause-specific hazard models for clinical covariates. Models were distinct for each site, and included all covariates shown. Rows are sorted by covariate and then estimated hazard ratio. For forest plots, the center box indicates estimated HR with whiskers showing the range of the 95% confidence interval for the HR. **b.** Exemplary clinical covariate association site specific cumulative incidence curves. Cumulative incidence curves of liver metastasis stratified by sex, lung metastasis stratified by median split age, bone metastasis stratified by tumor mutational burden quartiles, and peritoneal metastasis stratified by tumor mutational burden quartiles. **c.** Associations of functional mutations in gene with site metastasis. Each box represents the model results for different metastatic sites. Each point represents the results from a penalized cause-specific hazard model with the incidence of the specific metastatic sites predicted by the functional mutation status of a gene. Total distinct metastatic sites was included in each model as a cumulative time dependent variable. Points colored orange are significant (Beta term P-Value < 0.05). **d** Select cumulative incidence curves highlighting site-specific associations. Patients are stratified by the functional status of the given gene. SDHR, p = subdistribution hazard ratio p-value.

We then tested whether functional mutations in genes were associated with a change in the risk of specific metastatic sites. Using a cause-specific hazard model with a time-dependent covariate of total metastatic sites, we analyzed gene mutations associated with site specific incidence of metastasis (Fig. 5c). To test robustness of our results, we tested a different survival model (subdistribution, Methods) and different choices of reference times (i.e. setting time zero to be the time of sequenced biopsy), and observed consistent results (Methods, Supp Fig. 7). We observed that loss-of-function mutations in *PTEN* were associated with increased incidence of brain metastasis (SDHR = 1.7, 95% CI [0.98-2.92], p = 0.057; CSHR = 1.74, 95% CI [1.01-3.0], p = 0.046; adjusted CSHR = 1.82, 95% CI [1.06-3.12], p = 0.044, Fig 5d), consistent with previous literature linking *PTEN* with brain metastasis in melanoma^22^. We also observed broad site-specific associations with loss-of-function mutations in *B2M*, with higher incidence of metastasis in LN (SDHR = 1.74, 95% CI [1.21-2.49], p < 0.01), liver (SDHR = 2.0, 95% CI [1.06-3.78], p < 0.05), and lung (SDHR = 1.8, 95% CI [1.34-2.4], p < 0.01, Fig 5d). Other associations of functional gene mutations with site-specific metastases are shown in Supp Fig. 8.

To validate these associations, we tested select significant results in an independent cohort of patients with cutaneous melanoma from a different institution (MSK, Methods), including recapitulation of associations of liver, LN, and lung metastasis with *B2M* (Supp Fig. 3c-d). Overall, by using longitudinal models of metastasis, we identified site-specific associations with both clinical features and genomic alterations.

### GNA11/GNAQ mutations confer risk of liver metastasis in patients with cutaneous melanoma

We observed that gain-of-function alterations in *GNA11* and its paralog *GNAQ* were associated with increased incidence of liver metastasis (SDHR = 2.94, 95% CI [1.29-6.72], p = 0.01; SDHR = 2.87, 95% CI [1.1-7.5], p = 0.03 respectively). We validated this association in the independent cohort of patients with cutaneous melanoma described in the previous section (MSK) (Fig. 6a-b). Since *GNA11* and *GNAQ* are known paralogs and since driver mutations in these genes were mutually exclusive among our patients (and in other cancers), we combined the two groups of mutated patients into a single subgroup. This combined *GNA11*/*GNAQ* subtype had a large increase in incidence of liver metastasis (CSHR = 3.1, 95% CI [1.44-6.59], p = 0.004; SDHR = 3.01, 95% CI [1.59-5.7], p < 0.001) (Fig. 6c).

**Figure 6.**
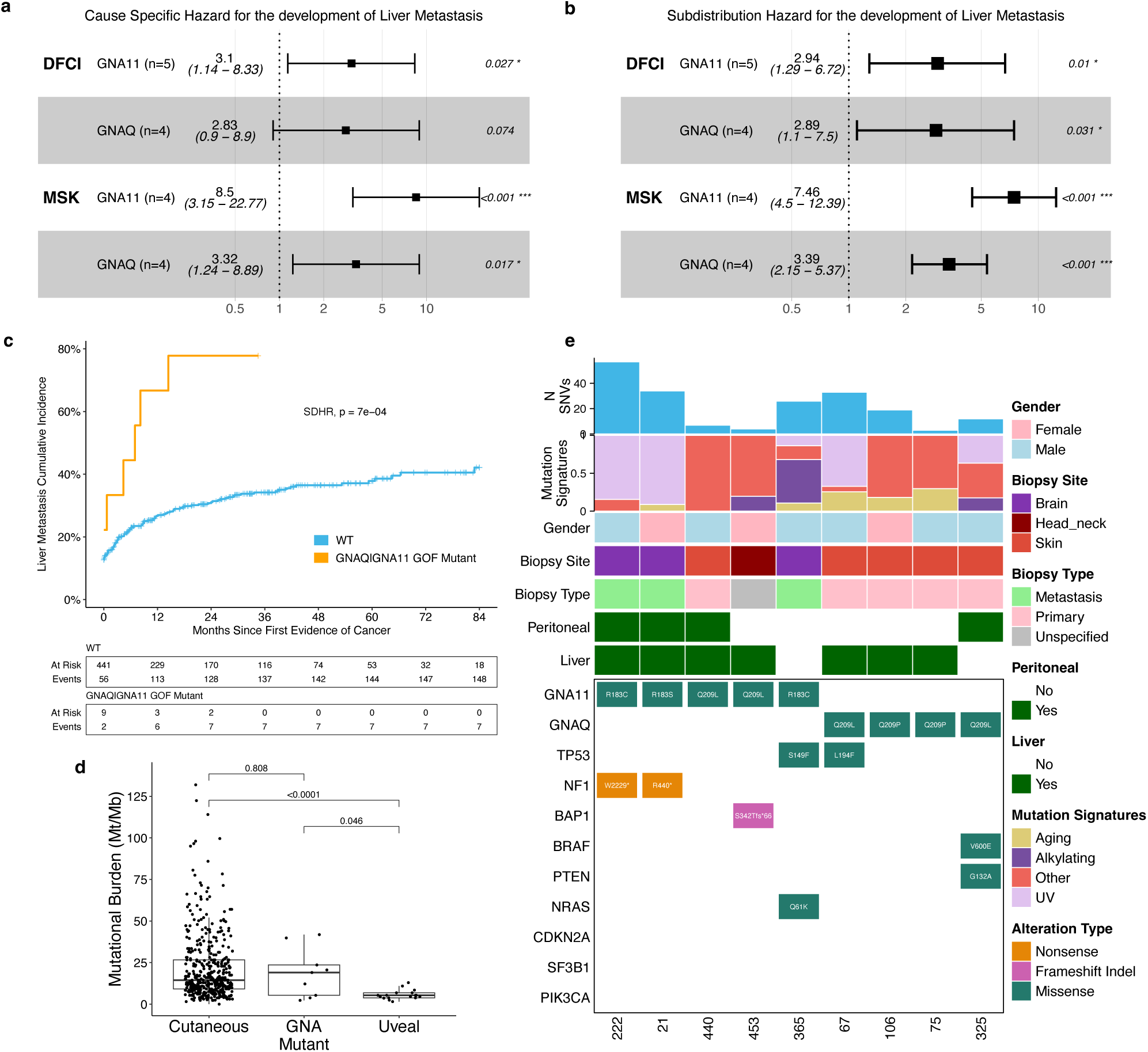
GNA11/GNAQ mutations confer liver metastasis in cutaneous melanoma patients. **a-b.** Computational validation of the GNA associations. Forest plot of (a) cause-specific or (b) subdistribution hazard models for incidence of liver metastasis stratified by *GNA11* or *GNAQ*. The first two rows show the results in our cohort, and the last two rows show the results of the same analysis in the MSK cohort. For forest plots, the center box indicates estimated HR with whiskers showing the range of the 95% confidence interval for the HR. **c.** Cumulative incidence curve of liver metastasis stratified by whether cutaneous patients had gain-of-function mutation in either *GNAQ* or *GNA11*. **d.** TMB comparison of GNA patients with other cutaneous patients in the cohort and uveal melanoma patients (N=17). Test bars represent the p values from wilcoxon rank sum tests. **e.** Comutation plot showing select clinical and genomic characteristics for all cutaneous patients in the cohort with either a *GNA11* or *GNAQ* functional mutation (N=9 patients). Each column represents a patient and sequenced biopsy. Mutational signature refers to the inferred relative contribution of UV-induced mutations, alkylating DNA damage, Aging effect, or other mutational signatures. Liver status refers to whether the patient had a liver metastasis in the follow-up period. SF3B1 had no functional mutations across the patients. Boxplots: Box limits indicate the IQR (25th to 75th percentiles), with the center line indicating the median. Whiskers show the values ranges up to 1.5 x IQR above the 75th or below the 25th percentiles.

Since uveal melanomas are driven by mutations in *GNA11* or *GNAQ* and canonically metastasize to the liver^23^, we hypothesized that *GNA11* and *GNAQ* mutations in cutaneous melanoma may confer a tumor-intrinsic propensity for liver tropism. To verify that our cutaneous melanomas with GNA-mutations were not misclassified uveal melanomas, we first reviewed the clinical history of these patients, confirming cutaneous melanoma diagnosis and no concurrent uveal melanoma. Second, these GNA11/GNAQ mutant cutaneous melanoma tumors had TMB levels consistent with cutaneous melanoma and significantly higher than uveal melanoma (Fig. 6d). Third, we characterized other clinical and genomic features of these patients (Fig. 6e). Of the 9 tumor samples, 5 were primary skin biopsies (consistent with cutaneous melanoma etiologies), 1 sample from head-neck, and 3 samples were brain biopsies. A UV mutational signature was detected in 5 of the 9 tumor biopsies. Of the 4 tumors without called UV signatures, 3 were characterized by a low number of SNVs which may have limited the ability to call the signature. Evaluating at the co-occurrence of other mutations, 2 patients had *NF1* mutations, 1 had a *BRAF* mutation, and 1 had an *NRAS* mutation, uncommon in uveal melanomas^23^. Other typical genes mutated in uveal melanoma such as *SF3B1*, *BAP1* were only seen in one of ten tumors. Lastly, four patients had a peritoneal metastasis in their disease history, a location less characteristic of uveal melanoma^24^.

Thus, clinical and genomic characterization confirmed that these tumors had clinical and genomic characteristics of cutaneous melanoma, and this, along with validation in an independent dataset, gives us confidence in our observation that cutaneous melanoma patients with *GNA11/GNAQ* mutations exhibit high rates of liver metastasis.

### Liver tropism of Gnaq/Gna11-driven cutaneous melanoma in genetically engineered mouse models

To examine the association between metastatic organotropism and driver mutations in pre-clinical models, we compiled the known alleles and sites of spontaneous metastases (Table 1) of previously described genetically engineered mouse (GEM) models of cutaneous melanoma. Specifically, we selected models where spontaneous sites of metastasis were assessed after the primary melanoma was initiated (following genetic activation and/or carcinogenic treatment) and the tumor was genomically characterized for driver mutations.

**Table 1.**
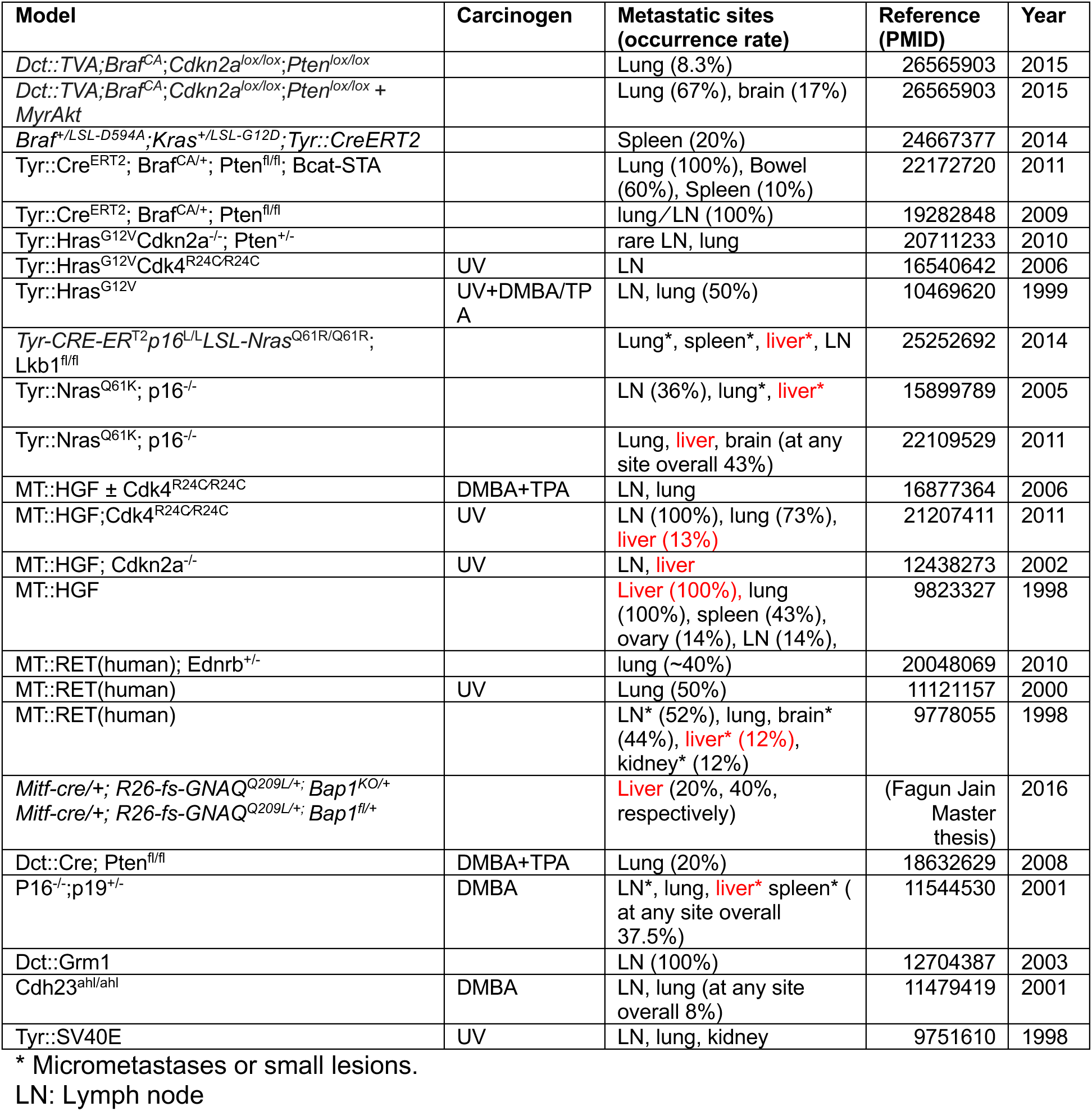
Spontaneous metastasis from the autochthonous cutaneous melanoma generated in the genetically engineered mouse models.

In these experimental model systems, whereas *Braf* and *Ret*-driven models were dominated by lung metastases, *Hras*-driven models with or without carcinogen treatment were dominated by lung and lymph node macrometastases with lesions at other sites mostly micrometastases. In contrast, liver metastases appeared more frequently in *Nras*- and *Gnaq*-driven models and *Hgf*-transgenic (tg) melanoma models, which frequently exhibited Q209 hotspot mutations in Gnaq/Gna11^25^. We have previously identified frequently mutated Q209 hotspots within *Gnaq/Gna11* in *Hgf*-transgenic melanoma^26^ which are consistent with the human data that *Gnaq/Gna11* mutations in melanoma are associated with liver metastasis.

We then evaluated the liver organotropism of a *Gna*-mutant melanoma (Fig. 7a) using C5, a GEM-derived allograft (GDA) line of cutaneous melanoma with spontaneous occuring *Gna11*^Q209L^ and *Trp53*^R172H^ mutations, generated from a DMBA-induced melanoma in a *Hgf-tg;Cdk4*^R24C^ C57BL/6 genetic background^26–28^ (Supp Fig. 9a). We labeled the C5 cells with firefly (ff) luciferase/H2B-GFP genes *ex vivo* (Methods) to enable bioluminescence imaging (BLI) for monitoring disease progression and then implanted into a syngeneic mouse for tumor expansion^29^. In a representative study, the grown tumor was then transplanted subcutaneously into five syngeneic hosts (Fig. 7b). After the primary tumors were resected upon reaching 500 mm^3^, mice were monitored by BLI for metastasis. Three of five mice developed macrometastases in liver, and one out of five had macrometastases in lung (Fig. 7c-d; confirmed by necropsy reports). Noticeably, the metastatic burden in liver was significantly higher than those in lung.

**Figure 7.**
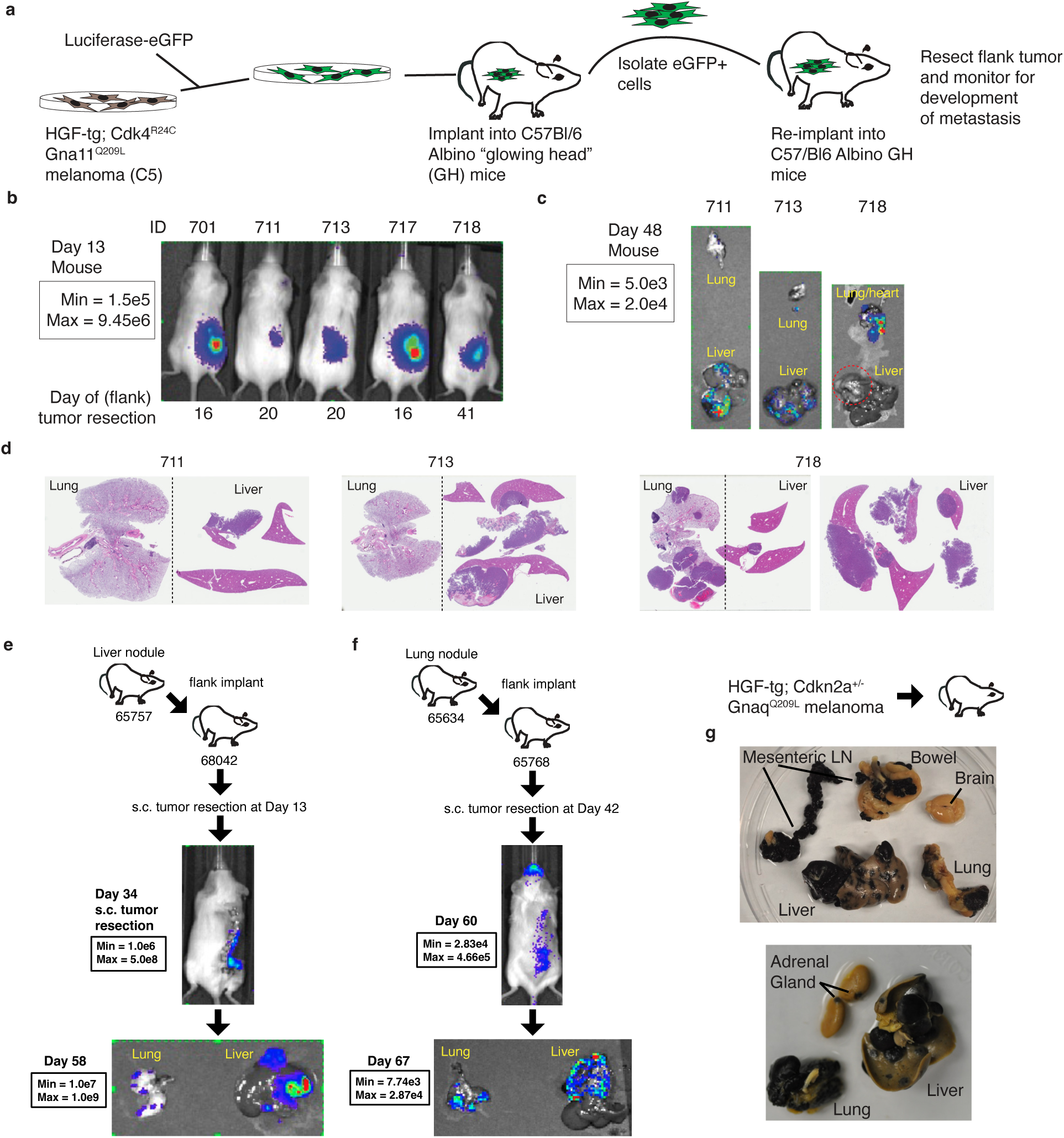
Liver metastasis in preclinical models of Gna11/Gnaq-mutatedm. **a.** Schematic of Gna11 melanoma model. Gna11-mutated HGF-tg;Cdk4^R24C^ melanoma line C5 was labeled with firefly (ff) luciferase/H2B-GFP genes ex vivo and then implanted into a syngeneic donor mouse. GFP+ cells were sorted from the harvested flank tumor or lung metastasis and then re-implanted into recipient albino C57Bl/6 GH mice. Primary flank tumors were resected and mice were monitored by bioluminescence imaging (BLI) for metastases. b. BLI of recipient mice on day 13 (n=5). Date below image represents day of primary tumor resection. c. BLI of harvested organs from recipients on day 48. d. Hematoxylin and eosin (H&E) staining of liver and lung from recipients on day 48. e. Exemplary experiment where liver metastases from a mouse with ff-Luc/GFP labeled GNA11 mutated tumors were re-implanted in the flank of recipient mice. Primary tumors were resected and mice monitored via BLI for metastases. BLI shown for mice on day 34. BLI of harvested organs from exemplary mouse at day 58 shown below. F. Exemplary experiment where lung metastases from a mouse with ff-Luc/GFP labeled GNA11 mutated tumors were re-implanted in the flank of recipient mice. Primary tumors were resected and mice monitored via BLI for metastases. BLI shown for mice on day 60. BLI of harvested organs from the mouse at day 67 shown below. g,h. Necropsy of organs harvested from two exemplary C57BL/6 mice implanted with HGf-tg;Cdkn2a^+/−^ derived, Gnaq^Q209L^-driven melanoma. Organs shown for mice include liver, lung, adrenal glands, brain, bowel, and mesenteric lymph nodes (LN).

In a parallel study, we tested whether tumors that had metastasized to one site could retain tropism to the liver. We harvested metastatic nodules from the lung or liver of the mice implanted with labeled C5 tumor, and then transplanted them subcutaneously into recipient mice, respectively, followed by monitoring metastasis by BLI after resection of the primary tumors. We observed that subcutaneous implants from liver metastases (Fig. 7e) and lung metastases (Fig. 7f) both resulted in similar metastatic phenotypes with high frequency metastases in the liver and lung and particularly high-burden metastases in the liver. Repeat experiments showing the robustness of this phenotype across implants from liver metastases, lung metastases, and subcutaneous primaries are summarized in Supp Fig. 9c-d, suggesting that the liver tropism is an intrinsic feature of this *Gna11*-mutant model.

Finally, we implanted a separate Gnaq^Q209L^-driven melanoma GDA recently derived from UV-irradiated Hgf-tg;Cdkn2a^+/−^ mice into syngeneic C57BL/6 mice and resected the primary tumors upon reaching 500 mm3 in a similar protocol (Methods). At the protocol-defined endpoint, the mice were euthanized for necropsy, and the organs were harvested for fixation. Across experiments, there was a large burden of liver metastases, high frequency of lung metastases, and variable metastases across other sites (Fig. 7g).

Taken together, these results demonstrated liver tropism of *Gna11/Gnaq*-mutated melanoma metastasis in immunocompetent mouse models and offers possible models by which to test further mechanisms or interventions.

## Discussion

In this study, we analyzed patterns of metastasis in clinico-genomic cohorts of cutaneous melanoma patients with longitudinal metastatic site annotations. We characterized multiple dimensions of metastasis, including overall metastatic potential and site-specific metastases, and provide the first (to our knowledge) large-scale analysis of patient-level patterns of metastasis, identifying major subgroups of patients with different patterns of metastasis differing by overall metastatic potential and site-dominant metastasis. We further identified novel genomic and clinical features associated with different rates of overall and site-specific metastases, as well as these different metastatic patient subgroups. We robustly validated these finding in independent patient cohorts and performed proof of principle validation in pre-clinical models of a genomic driver not previously associated with liver metastasis in cutaneous melanoma.

Among clinical factors, we observed that female sex was associated with decreased metastasis overall, which has been previously described in melanoma^30^ and NSCLC^31^. We observed the strongest protective effect of female sex in liver, peritoneal, and bone metastasis. While age was not associated with overall metastatic burden in our melanoma cohort (discordant with previous observations in NSCLC^31^), we observed a strong effect of age on increased rates of metastases to the lung and adrenal sites, and the highest age in the Lung-avid cluster of patients. Interestingly, recent work has found that aged lung microenvironments may induce reactivation of dormant melanoma cells and provide a permissive niche for metastasis^32^.

We observed a strong association between higher tumor mutational burden and fewer overall sites of metastasis consistent across all site-specific contexts. In our cluster analysis patients in the HM cluster had the lowest TMB. Interestingly, this was not observed in a prior multi-histology analysis, though there were trends in both directions in different tumor histologies^17^. In the context of treatment within cutaneous melanoma, higher TMB has been associated with response to immune checkpoint inhibition and longer overall and progression free survival^33^, potentially suggesting the role of immune activation and surveillance in limiting broad metastatic potential.

Indeed, a striking observation is that the two strongest driver mutations associations with high metastatic potential are *PTEN* and *B2M* loss-of-function mutations. *PTEN* mutations have been found to be associated with immunosuppressive microenvironments across different histologies, mediated through multiple mechanisms including regulation of immune cell trafficking (VEGFA expression) and autophagy^34–36^. *B2M* is required for MHC-I antigen presentation to T cells and acquired loss-of-function mutations are associated with development of resistance to immune checkpoint blockade^37,38^. Together, these observations suggest that immune surveillance mediated at least in part by T cells limits metastasis broadly across organ sites.

Importantly, we identify and independently validate global patterns of clinical metastasis, with key clinical and biological insights. Patient metastatic clusters are driven by two axes: overall metastatic potential (i.e. low vs high number of metastatic sites), and organotropism to specific sites (brain, lung, liver). These clusters have distinct clinico-genomic profiles, consistent with the hypothesis that there are distinct biological mechanisms leading to these different metastatic phenotypes. Specifically, we observed that patients with brain metastases in different clusters had distinct genomic and clinical features; the patients in the brain-tropic cluster were older and their tumors had almost twice the number of mutations compared to patients in the highly metastatic cluster. Further, while *PTEN* loss-of-function mutations have been previously shown to associate with brain metastases^17,22,39^ (and recapitulated in our cohort (Fig. 5c)), they strongly associated with increased overall metastatic potential (Fig. 3d). Consistent with this, they were enriched in the patient cluster defined by high metastatic potential rather than the brain-avid cluster (Fig. 4g), suggesting that the mechanism leading to brain metastasis in *PTEN* mutated melanomas is not unique to the brain microenvironment and leads to metastases at other sites. Taken together, our observations strongly suggest that site-specific metastasis can be driven by different biological mechanisms which manifest in distinct metastatic patterns.

We find a strong novel association of *GNAQ/GNA11* gain-of-function mutations with rapid development of liver metastases in this cutaneous melanoma cohort and validated this association in an independent clinical cohort. Prior literature has described a subset of cutaneous melanomas with *GNA11/Q* driver mutations^40^, and multiple lines of evidence (TMB, co-occurring driver mutations, UV mutation signatures, and clinical review) gives us confidence that these are indeed cutaneous melanomas.

Interestingly, *GNAQ/GNA11* mutations drive uveal melanomas (melanomas that arise in the eye), and when these tumors metastasize, they canonically metastasize to the liver. This suggests that *GNA11/Q* mutations may be associated with a tumor-intrinsic biological state with tropism to the liver. In preclinical models, liver is a rare metastatic site; however, in existing autochthonous (spontaneous) cutaneous melanoma mouse models, liver metastasis was enriched in *Gna11/q* mutant melanomas (Table 1). We further confirm striking and reproducible liver metastasis after flank injection (and survival surgery) of *Gna11*- and *Gnaq*-mutant cutaneous melanoma mouse models.

Other mechanisms of liver metastatic organotropism have been demonstrated, including PI3K/AKT signaling pathways identified using CRISPR screens, with *PIP4K2C* loss as a metabolic adaptation for metastatic colonization^7^. The mechanism of *GNAQ/11* mutations as drivers of liver organotropism are unclear. It was recently observed that *GNAQ*^Q209L^ mutations led to constitutive phosphorylation of MET (receptor for hepatic growth factor, HGF) in an immortalized melanocyte cell line^41^, and prior work has implicated the cMet/HGF axis in liver metastasis^42^, but whether this is the primary mechanism or whether other pathways co-regulate the process requires further mechanistic studies. Further, non-cutaneous melanomas with GNA-mutations (uveal, CNS^43^, and mucosal melanomas^44^) have poor treatment options and prognosis after metastasis. Manual review of two patients with rare CNS primary melanomas (excluded from the primary analysis) and *GNA11/GNAQ* mutations revealed that both ultimately developed liver metastasis, highlighting the potential broader impact of elucidating the relevant mechanisms (and potential therapeutic targets) across multiple hard-to-treat diseases.

Several limitations exist for this study. A fundamental limitation is that genomic sequencing was obtained from a single biopsy in the vast majority of patients; an implicit assumption in our analyses is that the genomic characteristics are consistent over time in the same patient, though multiple prior studies of matched tumor samples have shown that driver mutations are largely maintained between tumors^45^. Despite the size of our cohort, the relevant group sizes decrease quickly when we consider specific covariates (e.g. functional gene mutation, treatment, metastatic cluster), and larger cohorts would improve our power to detect these types of relevant effects. Differences in treatment were not explored in this study but certainly may impact metastatic outcomes or tumor evolution. Our validation in independent cohorts lends confidence for specific genomic findings, but we expect additional analysis of other large independent cohorts will yield additional discoveries, in addition to functional studies to further characterize the biological mechanisms underpinning our observations.

Taken together, our study presents a systematic, integrative, and novel analysis of clinical organotropism in cutaneous melanoma, providing insight into the patterns and drivers of metastasis. We nominate novel mechanisms of site-specific metastasis, laying the foundation for further studies to dissect these mechanisms and identify targets to interrupt the metastatic process. Broadly, our study provides a framework for deriving biological insights about the heterogeneity of metastasis in patients that is translatable to other cancer histologies.

## Methods

### Patient cohort, Processing, QC, Manual

We identified 881 patients with melanoma tumor NGS panel data generated between 2005-2022 and whose imaging reports were annotated by the NLP model. Using the model results from Kehl 2021^15^, we assembled a dataset of 26,117 artificial intelligence-annotated text imaging reports from 881 patients. Annotated imaging reports were filtered according to multiple criteria. Model predictions included presence of cancer, progression/response, worsening/improvement, and the presence of various metastatic sites. Possible sites of metastasis that were annotated included brain, bone, lymph node, lung, liver, adrenal gland, and peritoneal. No distinction was made between distant and regional lymph node metastasis. All patients were evaluated at the Dana-Farber Cancer Institute, and each patient had at least one tumor biopsy sequenced with OncoPanel, a next-generation DNA sequencing panel that identifies mutations in cancer-related genes^18^. Biopsy site of sequencing varied by patient, and no requirement was made for cohort inclusion. The melanoma cohort was limited to those patients with cutaneous melanoma (N=603). Patients with melanoma and an unknown primary site were included as cutaneous if they had evidence of cosmic^46^ mutational signature 7 (ultraviolet exposure) (N=142, filtered to N=74). Upon manual review, we removed one patient due to having a meningeal melanoma and removed another patient for having a spinal melanoma. Multiple criteria were used for both patient and scan inclusion. Scans were limited to those that were annotated as having a presence of cancer by the NLP model, along with being an MRI, CT, or PET scan. This was done to limit annotations to more accurate and task-specific scans. Patients were included if they had at least one scan that passed these criteria, thus having radiographic evidence of cancer. This resulted in a total of 598 patients with cutaneous melanoma. Lastly, we limited our cohort to patients who developed metastatic cutaneous melanoma by examining the disease extent of patients at first radiographic evidence of cancer. Patients who did not have metastatic disease at this point, and did not have a metastatic site predicted in the follow-up period, were excluded for not having sufficient evidence of ever having metastatic melanoma. We implemented a time cutoff of 84 months to limit timespan of metastatic development to relevant follow-up periods, retaining the full follow-up for over 90% of patients. No minimum follow-up was required for cohort inclusion. For quality control, we evaluated the performance of the NLP metastatic annotations on a subset of 66 melanoma patients with manual annotations from radiographic reports within 4 months prior to systemic immunotherapy treatment. Within this time range, the NLP model performed well across the metastatic sites, with median sensitivity and specificity of 0.88 and 0.94, respectively (Supp Fig. 2a-b).

### DNA sequencing and Functional Annotation

All patients included in the cohort had OncoPanel targeted panel sequencing for at least one melanoma biopsy, allowing identification of tumor somatic alterations^18^. These samples consisted of three versions of OncoPanel Sequencing, with the number of sequenced genes differing between them. Version 1 sequenced 275 genes (n=19), version 2 sequenced 300 genes (n=79), and version 3/3.1 sequenced 447 genes (n=374). To ensure fair comparison across samples, we limited to only genes that were shared across all versions on OncoPanel. This led to 239 genes of interest. Gene level somatic mutations were filtered for functional impact using the OncoKB database^47^. Alterations annotated with oncogenic levels of “Likely Oncogenic”, “Oncogenic”, “Resistance” or annotated with mutation effect of “Gain-of-function”, “Likely Gain-of-function”, “Likely Loss-of-function”, “Likely Switch-of-function”, “Loss-of-function”, “Switch-of-function” were included as being functionally relevant. After gene level filtering, 158 genes still had at least 1 functional alteration across the cohort. When patients had more than one sequenced sample (17 patients), mutation status for genes were aggregated across all sequenced biopsies. Thus, mutation status in a gene indicates whether a patient had a functionally relevant mutation in that gene across any of their available sequenced biopsies. Tumor mutational burden estimates were also averaged across all samples for patients with multiple samples. Finally, genes included in statistical testing were limited to those that were altered in at least 3 patients across the cohort, leading to 96 genes analyzed in this study. For more interpretable testing of genomic associations, we categorized alterations in genes as loss-of-function (LOF), gain-of-function (GOF), or other based on the OnkoKB annotation details.

For mutation signature calling, we used *deconstructSigs* (v1.8.0). Signatures were from the single base substitution signatures found in the Catalog of Somatic Mutations in Cancer (COSMIC) database^46^.

### Survival modeling

Time to event analysis was set up as follows: Start time was defined as the date of the first scan predicted to have evidence of cancer based on NLP model annotations of imaging reports. Time of metastasis was defined as the date of the first imaging report predicted positive for each site. Patients who did not have a site in the follow-up time were censored, and those who died were marked for the competing event. For overall survival estimation, we used the Kaplan-Meier method, and to model the effect of covariates on survival we used a Cox proportional hazard model. We used cause-specific hazard and subdistribution hazard models to account for competing events and determine the effect of covariates on metastasis. Cause-specific models were run in R using *coxph* from the *survival* (v3.5-7) package, with death being marked as a censoring event. Penalized cause-specific models using Firth’s penalized likelihood for Cox regression were run in R using *coxphf* from the *coxphf* (v1.13.4) package, with death again marked as a censoring event. The penalized model was used to account for cases of complete separation. Subdistribution hazard models were run in R using *crr* from the *tidycmprsk* (v0.2.0) package, with death coded as a competing event. For estimation of mean cumulative functions, we used the *mcf* function from the *reda* (v0.5.4) package. We used Andersen-Gill models for modeling new metastatic sites as recurrent events. Andersen-Gill models were run in R using *coxph* with the general format: *coxph(Surv(tstart,tstop,metastasis)∼covariate, cluster=ID, …)*. Time dependent covariates of either total metastatic sites or site-specific incidence were created using the *tmerge* function from the *survival* package and were included as mentioned for certain models.

### Unsupervised clustering and assessment

Bernoulli mixture models were estimated using the R *flexmix* package (v2.3-19)^48^. We clustered patients by their status for metastatic sites in the follow-up period. For each value for k components (ranging from 1-7), we ran 1,000 random initializations for 500 maximum iterations. The expectation maximization would stop when the change of log-likelihood was smaller than 1×10^−6^ and kept the model with the maximum likelihood. We calculated Akaike information criterion (AIC) and silhouette scores for each value of k components, noting that the minimum AIC was at 4 mixture components, with similar results for neighboring values (Supp Fig. 4a), while there was a large increase in median silhouette score from 4 to 5 components (Supp Fig. 4b-c). Due to this substantial increase in median silhouette score with a minimal change in AIC, we decided to use k=5 components for the final Bernoulli mixture model.

To test how the estimated clusters compared relative to a random dataset with no association between metastatic sites, we generated 1000 random site occurrence datasets by permuting our observed site data to create a null distribution of median silhouette scores for a k=5 Bernoulli mixture model. Using this null distribution, we generated significant empirical p-values for the metric from our observed data (Supp Fig. 4d).

To further explore how the structure of the clusters fluctuated with different component numbers, we plotted the frequency of metastatic sites across clusters for the optimal model at each value of k (Supp Fig. 4e). We further created a cluster tree showing the movement of patients between clusters at different numbers of components (Supp Fig. 4f), observing clear presence of similar cluster characteristics at both lower and higher numbers of components, but a low degree of inter-cluster movement past 5 components.

To test for enrichment of mutations in any cluster, we performed Fisher exact tests for whether a patient’s tumor had a functional mutation in each gene with whether a patient is in one cluster vs any other.

To classify patients into clusters, we calculated the posterior class probabilities for each cluster by using the baseline frequency of each cluster as the prior probability and using site frequencies to calculate the likelihood of observing those metastatic sites given a cluster^48^. To account for cases where the site frequency of a cluster is 0, a weight is added to make the posterior probabilities defined. We then assign patients to the cluster with the maximum posterior probability.

### Validation Cohorts

We utilized two validation cohorts of patients with cutaneous melanoma.

For our cluster validation cohort, we utilized the nationwide, longitudinal Flatiron Health database, an electronic health record-derived database comprising deidentified patient-level structured and unstructured data, curated via technology-enabled abstraction from approximately 280 cancer clinics (∼800 sites of care) in the United States^49,50^. The majority of patients in the database originate from community oncology settings; relative community/academic proportions may vary depending on study cohort. The data are deidentified and subject to obligations to prevent reidentification and protect patient confidentiality. Institutional review board approval was obtained prior to conducting the study (IRB#2000036075). Patients with metastatic melanoma treated in the front line setting with immunotherapy (ipilimumab + nivolumab, anti-PD1 monotherapy, or nivolumab + relatlimab) or BRAF/MEK inhibitors were included (n=5,124). Patients without documented metastatic sites or overall survival data were excluded.

For genomic validation, we utilized a cohort of 1,801 patients with cutaneous melanoma with MSK-IMPACT targeted tumor sequencing^51^ and NLP derived longitudinal metastatic annotations for all sites of interest except peritoneum^16^.

### Mouse models of metastatic Gnaq/Gna11-mutated cutaneous melanoma

GEM-derived allograft (GDA), C5, was generated from a Hgf-tg;Cdk4^R24C^ C57BL/6 mouse following DMBA treatment, as described previously^26^. Additional mutations were characterized by exome sequencing, including Gna11^Q209L^ (Supp Fig. 9b). The generation of firefly (ff) Luc-eGFP-labeled C5 GDA has been described previously^29^. Briefly, the C5 GDA was transplanted into syngeneic C57BL/6 mice subcutaneously. Harvested C5 tumors from the transplanted hosts, mixed with culture medium containing lentivirus of FerH-ffLuc-IRES-H2B-GFP (IRES, internal ribosome entry sequence; H2B-GFP, Histone 2B-tagged green fluorescence protein). Following transduction by spinoculation^52^, the medium was removed, and the cells were subcutaneously transplanted into albino C57BL/6^cbrd/cbrd^ glowing head (cbrd GH) mice pre-tolerized with ffLuc and GFP^29^.

To test the metastatic potential, ff Luc/H2B-GFP-labeled C5 tumors were transplanted into cbrd GH mice subcutaneously. Once the primary tumors reached a size of 500mm3, tumors were surgically resected and mice monitored via bioluminescence (BL) using the Xenogen IVIS system^52^. At the endpoint defined by the sickness behavior and moribund status, the mice were euthanized for necropsy, and the organs were harvested for BLI imaging ex vivo.

In another GEM model, melanoma was induced in Hgf-tg;Cdkn2a^+/−^ mice by UV irradiation at neonatal day 3. The procedure was described previously except that the mouse strain background used in this study is C57BL/6^53^. One of the melanoma tumors was harvested from a donor mouse and transplanted into syngeneic recipient for expansion and further passage as GDA (Fig. 7g). The endpoint was also defined by the sickness behavior and moribund status.

### Statistical analysis and visualization

All statistical analyses were performed in R (v4.2.1 “Funny Looking Kid”). For the comparison between continuous clinical and molecular features, the Mann-Whitney test was used. For association of binary variables, Fisher’s exact test was used. Reported p-values represent nominal p-values unless otherwise stated. All statistical tests performed were two-sided unless otherwise stated.

Visualizations were created with the R packages *ggplot2* (v3.4.2), *cowplot* (v1.1.1), *ggpubr* (v0.6.0), *ComplexHeatmap* (v2.14.0), *survminer* (v0.4.9), and *tidycmprsk* (v0.2.0). The Adobe Illustrator software was used to arrange figures.

## Declarations

### Funding

This research was supported by the Intramural Research Program of the National Cancer Institute (C.P.D.), the National Institute of Health (K08 CA234458, U2CCA233195; D.L.), and the Doris Duke Clinical Scientist Development Award (D.L.).

### Conflict of interest/Competing interest

R.H. reports advisory/consulting with Immunocore and research funding from Pfizer and Novartis. K.L.K reports research funding from Meta. J.J.I. reports the following relationships: personal fees from Related Sciences (consulting), Danger Bio (consulting/equity), Phenomic AI (consulting / equity), Kronos Bio (equity) and grants from AstraZeneca (sponsored research agreement) outside of the submitted work. A.N.S. reports advisory board / personal fees with Bristol-Myers Squibb, Immunocore, Novartis, and Erasca, along with trial support to institution from Bristol-Myers Squibb, Immunocore, Novartis, Targovax, Pfizer, Alkermes (Mural Oncology), Checkmate Pharmaceuticals, Foghorn Therapeutics, Linnaeus Therapeutics, Lovance Biotherapeutics, Obsidian Therapeutics. G.M.B. has sponsored research agreements through her institution with: Olink Proteomics, Teiko Bio, InterVenn Biosciences, Palleon Pharmaceuticals. She served on advisory boards for: Iovance, Merck, Moderna, Nektar Therapeutics, Novartis, Replimune, and Ankyra Therapeutics. She consults for: Merck, InterVenn Biosciences, Iovance, and Ankyra Therapeutics. She holds equity in Ankyra Therapeutics. F.S.H reports grants and personal fees from Bristol-Myers Squibb and Novartis, personal fees from Merck, Compass Therapeutics, Apricity, Bicara, Checkpoint Therapeutics, Genentech/Roche, Bioentre, Gossamer, Iovance, Catalym, Immunocore, Kairos, Rheos, Bayer, Zumutor, Corner Therapeuitcs, Puretech, Curis, Astra Zeneca, Pliant, Solu Therapeutics, Vir Biotechnology, and 92Bio, outside the submitted work; In addition, Dr. Hodi has a patent Methods for Treating MICA-Related Disorders (#20100111973) with royalties paid, a patent Tumor antigens and uses thereof (#7250291) issued, a patent Angiopoiten-2 Biomarkers Predictive of Anti-immune checkpoint response (#20170248603) pending, a patent Compositions and Methods for Identification, Assessment, Prevention, and Treatment of Melanoma using PD-L1 Isoforms (#20160340407) pending, a patent Therapeutic peptides (#20160046716) pending, a patent Therapeutic Peptides (#20140004112) pending, a patent Therapeutic Peptides (#20170022275) pending, a patent Therapeutic Peptides (#20170008962) pending, a patent THERAPEUTIC PEPTIDES Therapeutic Peptides Patent number: 9402905 issued, a patent METHODS OF USING PEMBROLIZUMAB AND TREBANANIB pending, a patent Vaccine compositions and methods for restoring NKG2D pathway function against cancers patent number: 10279021 with royalties paid, a patent antibodies that bind to MHC class I polypeptide-related sequence A patent number: 10106611 issued, a patent ANTI-GALECTIN ANTIBODY BIOMARKERS PREDICTIVE OF ANTI-IMMUNE CHECKPOINT AND ANTI-ANGIOGENESIS RESPONSES Publication number: 20170343552 pending, and a patent Antibodies against EDIL3 and methods of use thereof pending.

All remaining authors have declared no conflicts of interest.

### Ethics approval

This retrospective study and associated informed consent procedures were approved by the central Ethics Committee (EC) of the Dana Farber Cancer Institute (IRB 20-684 and IRB 16-360). Approval by the local EC was obtained by investigators if required by local regulations.

All mouse experiments were performed in accordance with Animal Study Protocols (11-044 and 11-007) approved by the Animal Care and Use Committee, NCI-Frederick, NIH. NCI is accredited by the Association for Assessment and Accreditation of Laboratory Animal Care and follows the Public Health Service Policy on the Care and Use of Laboratory Animals. All animals used in this research project were cared for and used humanely according to the following policies: the US Public Health Service Policy on Humane Care and Use of Animals; the Guide for the Care and Use of Laboratory Animals (2011); and the US Government Principles for Utilization and Care of Vertebrate Animals Used in Testing, Research, and Training (1985).

### Material availability

Further information and requests for resources and reagents should be directed to and will be fulfilled by D.Liu as corresponding author. Requests for reagents derived from mouse models (alleles, tumor tissue, cell lines) developed in this study will require a material transfer agreement as per NIH guidelines.

### Code availability

Software environments and code to reproduce all analysis and generate figure plots is available at (https://github.com/davidliu-lab/melanoma_organotropism_manuscript). Additional reasonable requests for code will be promptly reviewed by the senior authors to verify whether the request is subject to any intellectual property or confidentiality obligations and shared to the extent permissible by these obligations.

### Data availability

All data necessary for figure generation are in supplementary tables or data available at (https://github.com/davidliu-lab/melanoma_organotropism_manuscript). Any reasonable requests for raw and analyzed data and materials will be promptly reviewed by the senior authors to determine whether the request is subject to any intellectual property or confidentiality obligations. Patient-related information not included in this paper may be subject to patient confidentiality.

### Authors’ contributions

T.J.A and D.Liu conceived and designed the overall study. C.D. performed mouse experiments and contributed manuscript sections. T.J.A., G.T. acquired and processed data, and/or performed data analysis. Z.W and R.E assisted with data acquisition. M.M., H.F., M.M.H., K.Khaddour, and C.F. assisted with clinical annotation and data acquisition. D.Lee, A.P., J.I. performed validation of clustering results and interpreted data. J.J., N.S., A.S., performed validation of genomic results. G.T., M.Glettig, and K.P.B. interpreted data. K.Kehl., M.Griffin, A.G., J.I., K.P.B., F.S.H., G.M.B., and R.H. contributed key discussions. T.J.A. and D.Liu interpreted the data and wrote the manuscript. All authors reviewed and edited the manuscript.

## Supplemental Information

**Supplemental Figure 1.**
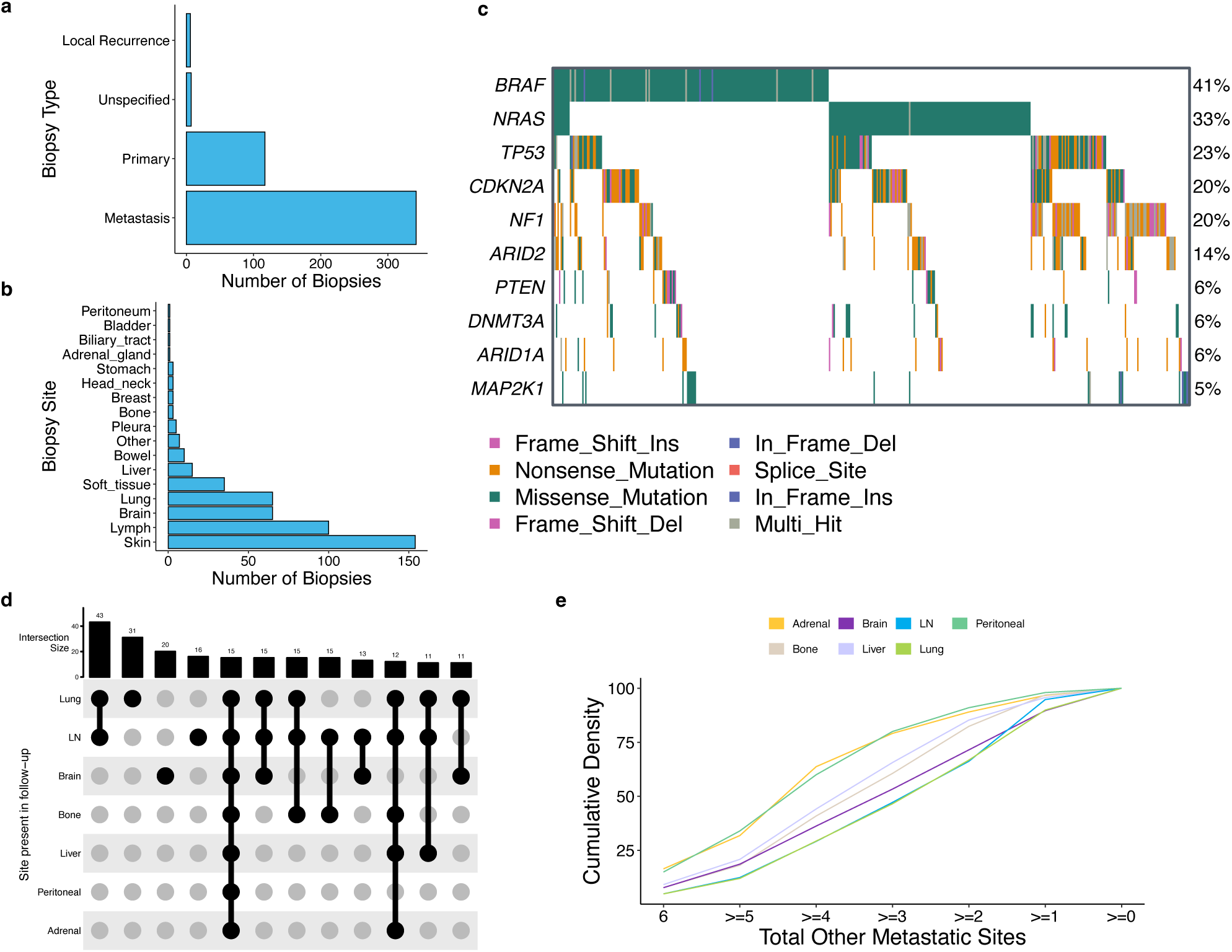
Additional cohort overview and site metastasis outcomes. **a.** Distribution of biopsy types across the sequenced samples. **b.** Distribution of biopsy sites across the sequenced samples. **c.** Overview of the ten most frequent alterations in the cutaneous melanoma patients. Rows are genes, and columns represent individual patients. Total frequency of alterations across the cohort for each gene are labelled on the right side. BRAF, NRAS, and NF1 frequencies are similar to expected genomics of melanoma. **d.** Upset plot showing the frequency of lifetime metastatic patterns across the cohort. Patterns that occur in at least 10 patients are shown. **e.** Cumulative density curves of total other metastatic sites stratified by whether patients had that site in lifetime.

**Supplemental Figure 2.**
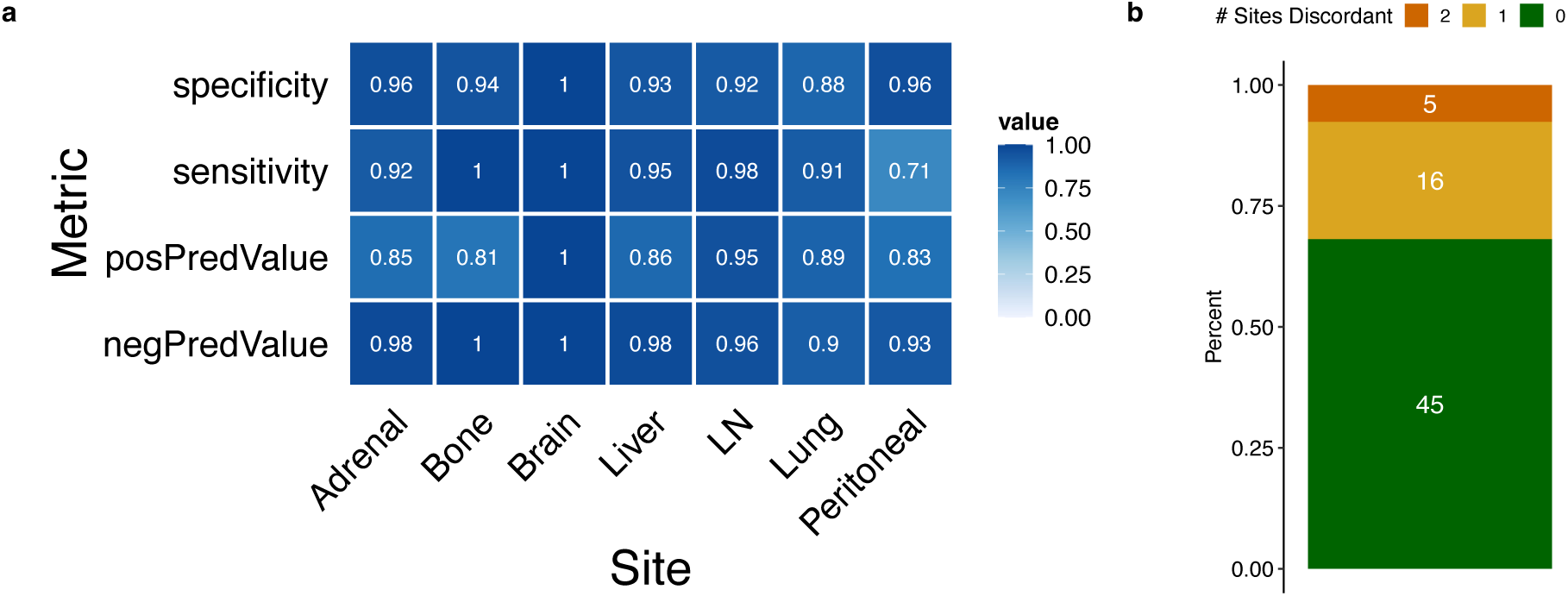
NLP annotation performance compared to manual annotations. **a.** Performance metrics for NLP annotations compared to manual annotation ground truth for each metastatic site. **b.** Stacked barplot showing the number of discordant sites by patient between the NLP annotations and manual annotations. A low number of patients have multiple discordant site labels.

**Supplemental Figure 3.**
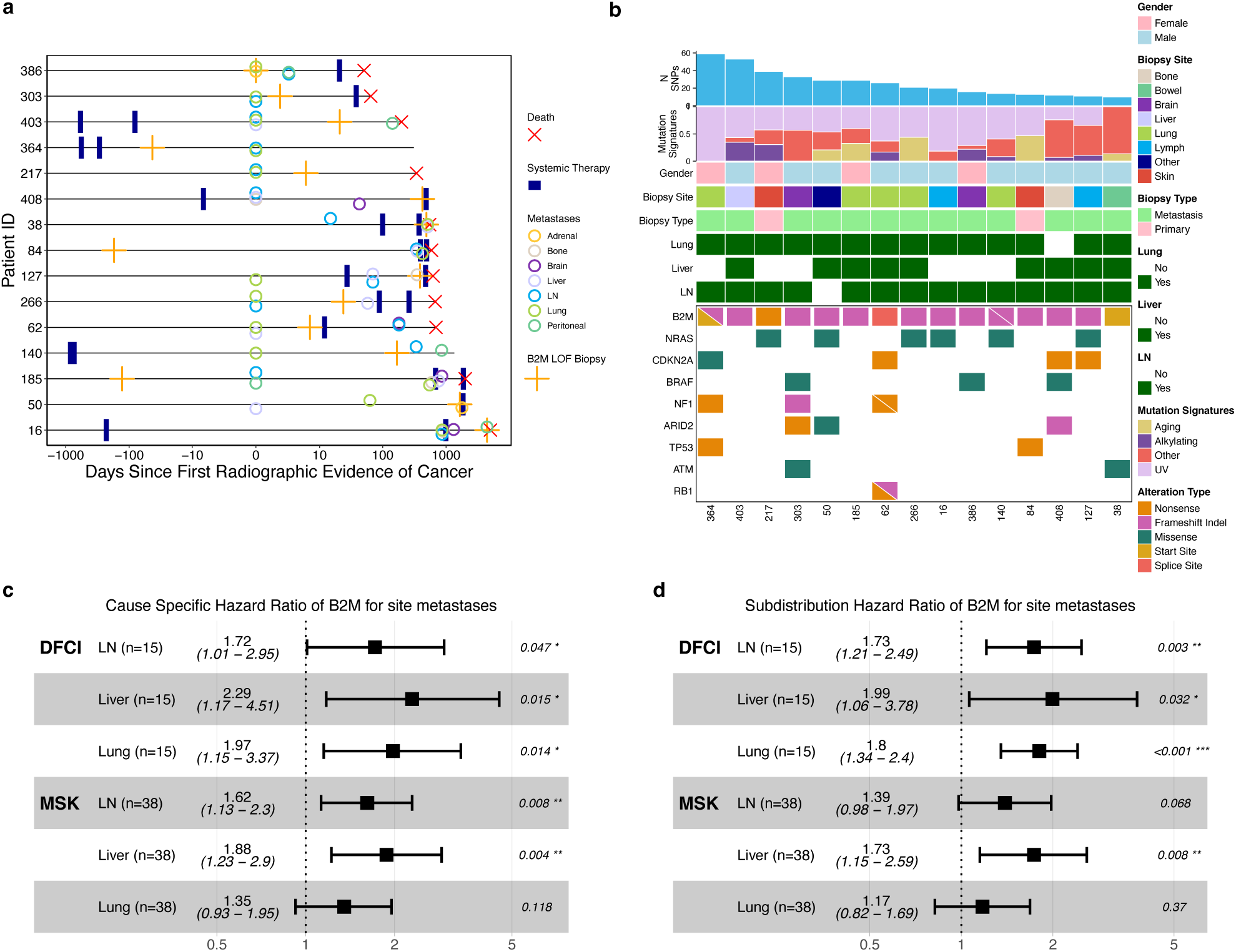
B2M Exploration and Validation. **a.** Visualization of the patient history for *B2M* mutant patients. Grey bars represent the follow-up time for the patients based on imaging reports, starting at first radiographic evidence of cancer. Red Xs note death if occurred, and orange crosses mark the time at which the *B2M* mutated biopsy was taken. Colored dots represent time of first metastasis at a site, derived from our NLP annotations. Width of treatment bars correspond to length of treatment. Treatment that didn’t have end date available were capped at 1 month after start time. **b.** Comutation plot showing clinical and genomic characteristics for all cutaneous patients in the cohort with *B2M* functional mutation (N=16 patients). Each column represents a patient and sequenced biopsy. Mutational signature refers to the inferred relative contribution of UV-induced mutations, alkylating DNA damage, Aging effect, or other mutational signatures. Liver, lung, LN status refers to whether the patient had that metastatic site in the follow-up period. **cd.** Computational validation of the B2M site metastasis associations. Forest plot of (c) cause-specific or (d) subdistribution hazard models for incidence of liver, lung, or LN metastasis stratified by *B2M*. The first three rows show the results in our cohort, and the last three rows show the results of the same analysis in the MSK cohort. For forest plots, the center box indicates estimated HR with whiskers showing the range of the 95% confidence interval for the HR.

**Supplemental Figure 5.**
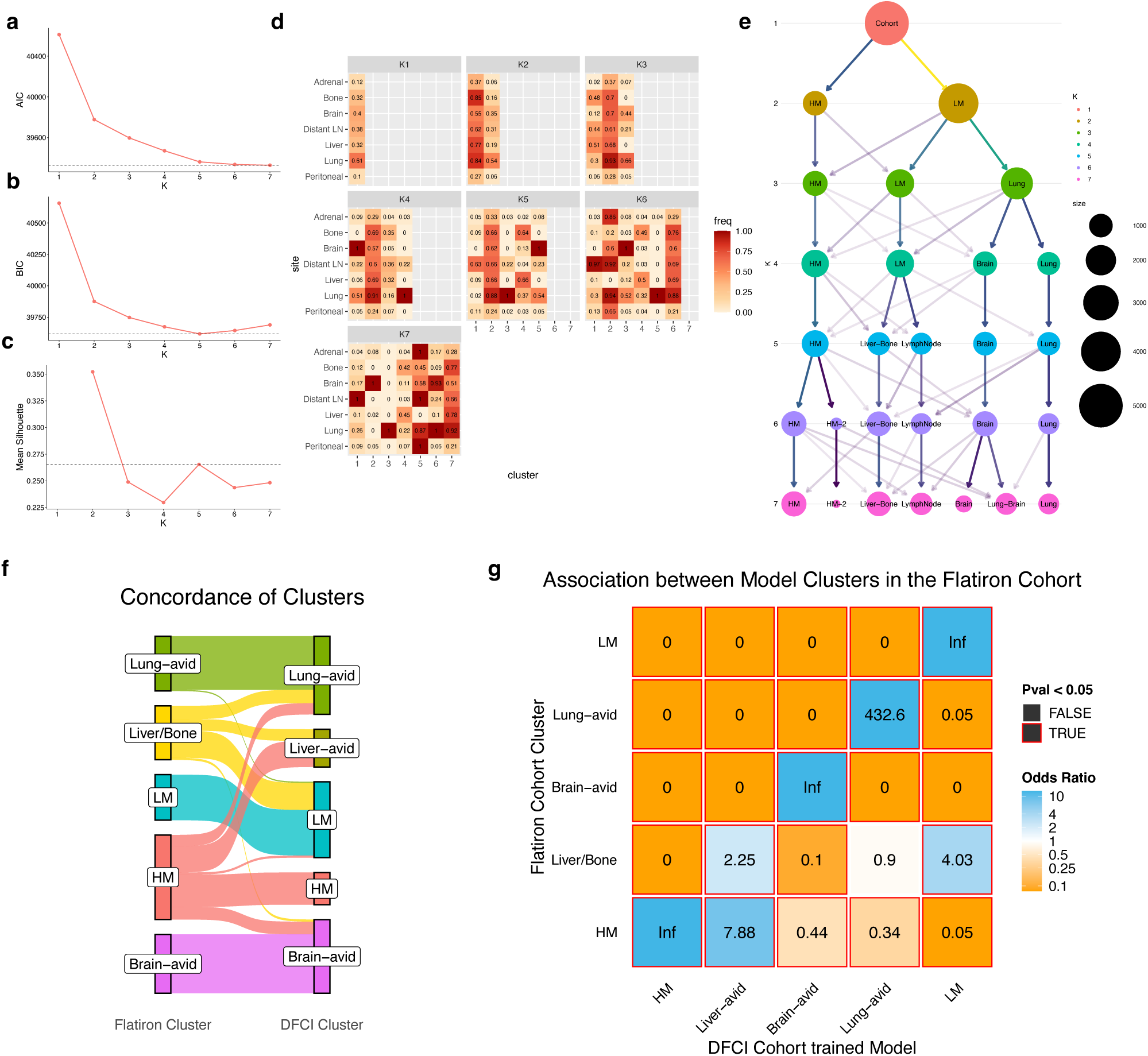
Flatiron cohort cluster validation. **a.** Plot of AIC metric for each value of k components for the Bernoulli mixture model. **b.** Plot of BIC metric for each value of k components for the Bernoulli mixture model. **c.** Plot of mean silhouette score for each value of k components for the Bernoulli mixture model. **d.** Heatmap of site frequency by cluster for each value of k components of the Bernoulli mixture model. **e.** Clustree visualization showing the size, interpretation, and movement of clusters as the value of k components changes in the Bernoulli mixture model. Darkness of arrows represent higher quantities of patients moving to a different cluster at different values of k components. Size of circles signify number of patients in the cluster. Color of circles signify value of k. **f.** Sankey Diagram showing the clusters of the Flatiron cohort when called with the DFCI cohort trained model. **g.** Associations between clusters of Flatiron cohort patients called with model derived from either the Flatiron or DFCI cohort. Each square represents a Fisher exact test odds ratio of whether a patient is classified in a specific cluster with the DFCI trained model vs a specific cluster with the Flatiron trained model. Inf represents cases where all patients put into the cluster in one model were put into that same cluster in the other model or the case where none of the patients who were not in that cluster in one model are moved into that cluster in the other model. Red borders represent tests with P-value < 0.05.

**Supplemental Figure 6.**
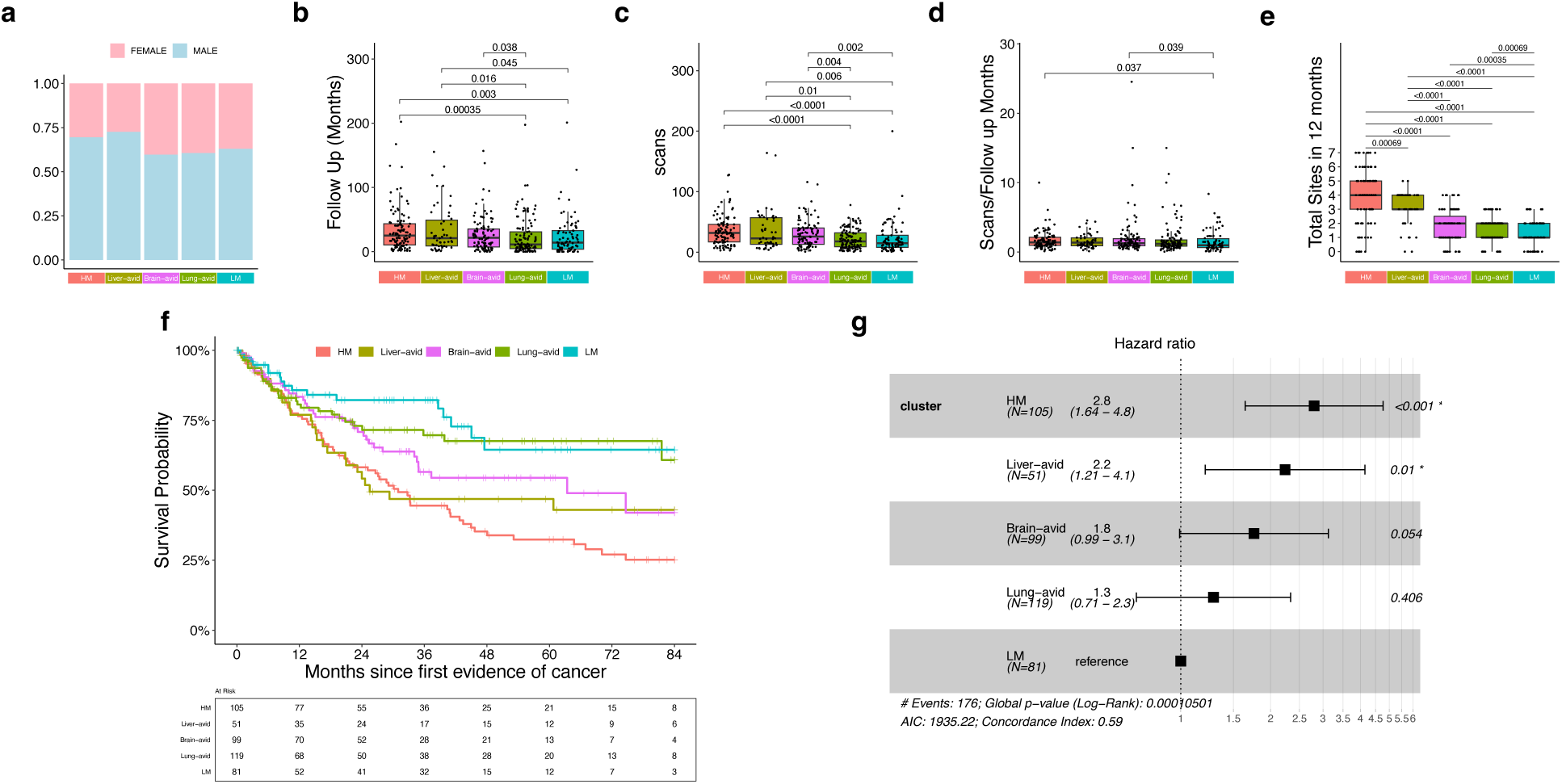
Addition analysis of cluster covariates. **a.** Stacked barplot of sex proportions by cluster. **b.** Boxplots of follow up time by cluster. **c.** Boxplots of scan number by cluster. **d.** Boxplots of scans per month of follow-up by cluster. **e.** Boxplots of total metastatic sites within the first 24 months of follow-up by cluster. **f.** Kaplan-Meier curves of overall survival stratified by cluster. **g.** Forest plot of cox proportional hazard model comparing survival by cluster. Reference group is the LM cluster. For forest plots, the center box indicates estimated HR with whiskers showing the range of the 95% confidence interval for the HR. Boxplots: Box limits indicate the IQR (25th to 75th percentiles), with the center line indicating the median. Whiskers show the values ranges up to 1.5 x IQR above the 75th or below the 25th percentiles. Comparison bars represent the results of wilcoxon tests. Only significant comparisons are shown.

**Supplemental Figure 7.**
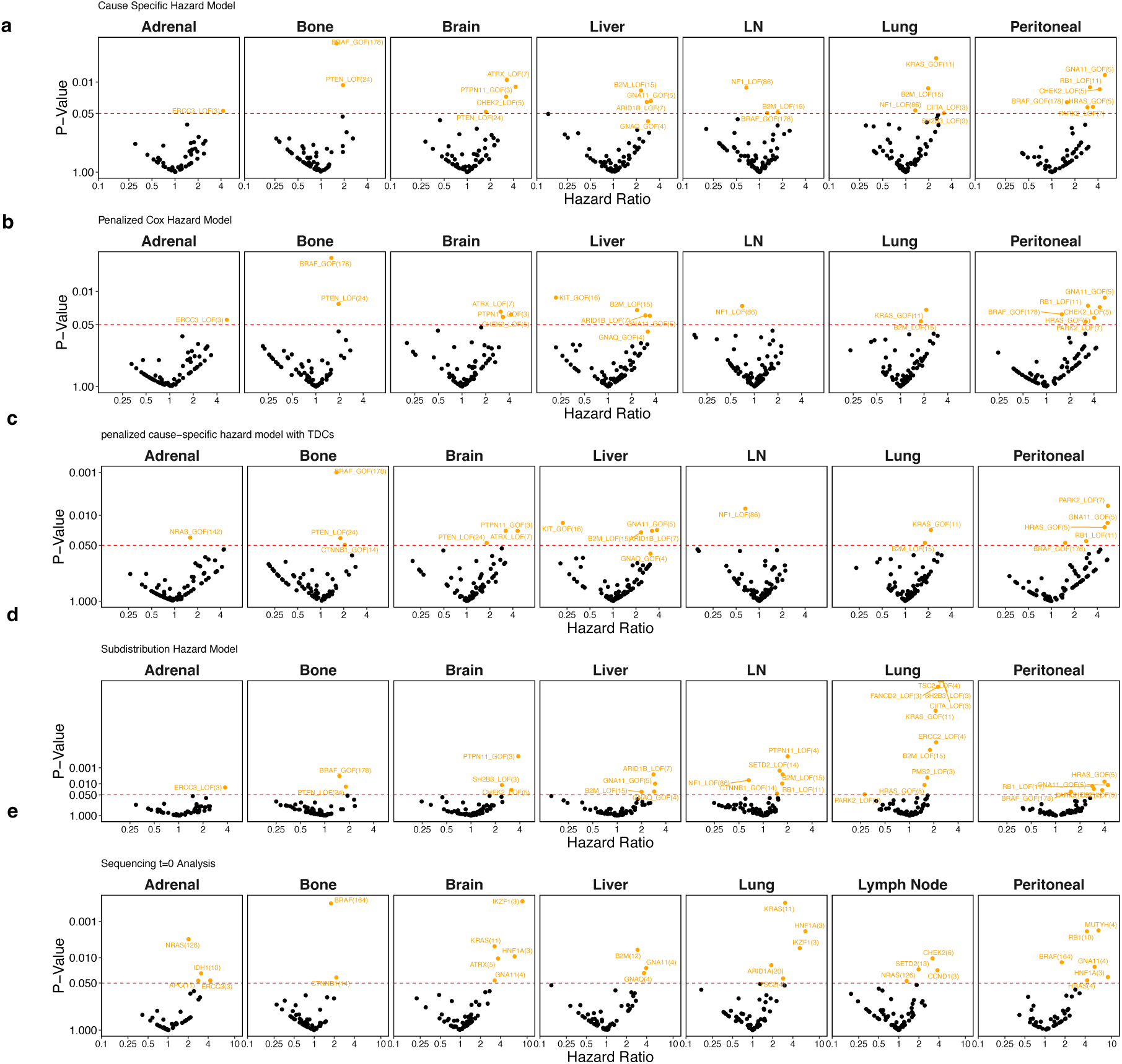
Alternate Time to Metastasis Models. Associations of functional mutations in gene with site metastasis. Each box represents the model results for different metastatic sites. Each point represents the results of a model with the incidence of the specific metastatic sites predicted by the functional mutation status of a gene. Points colored are significant (Beta term p< 0.05) **a.** Volcano plots of cause-specific hazard model results. **b.** Volcano plots of subdistribution hazard model results. **c.** Volcano plots of penalized cause-specific hazard model results. **d.** Volcano plots of penalized cause-specific hazard model with total distinct metastatic sites was included as a cumulative time dependent variable. **e.** Volcano plots of sequencing time 0 based data with cause-specific hazard model results.

**Supplemental Figure 8.**
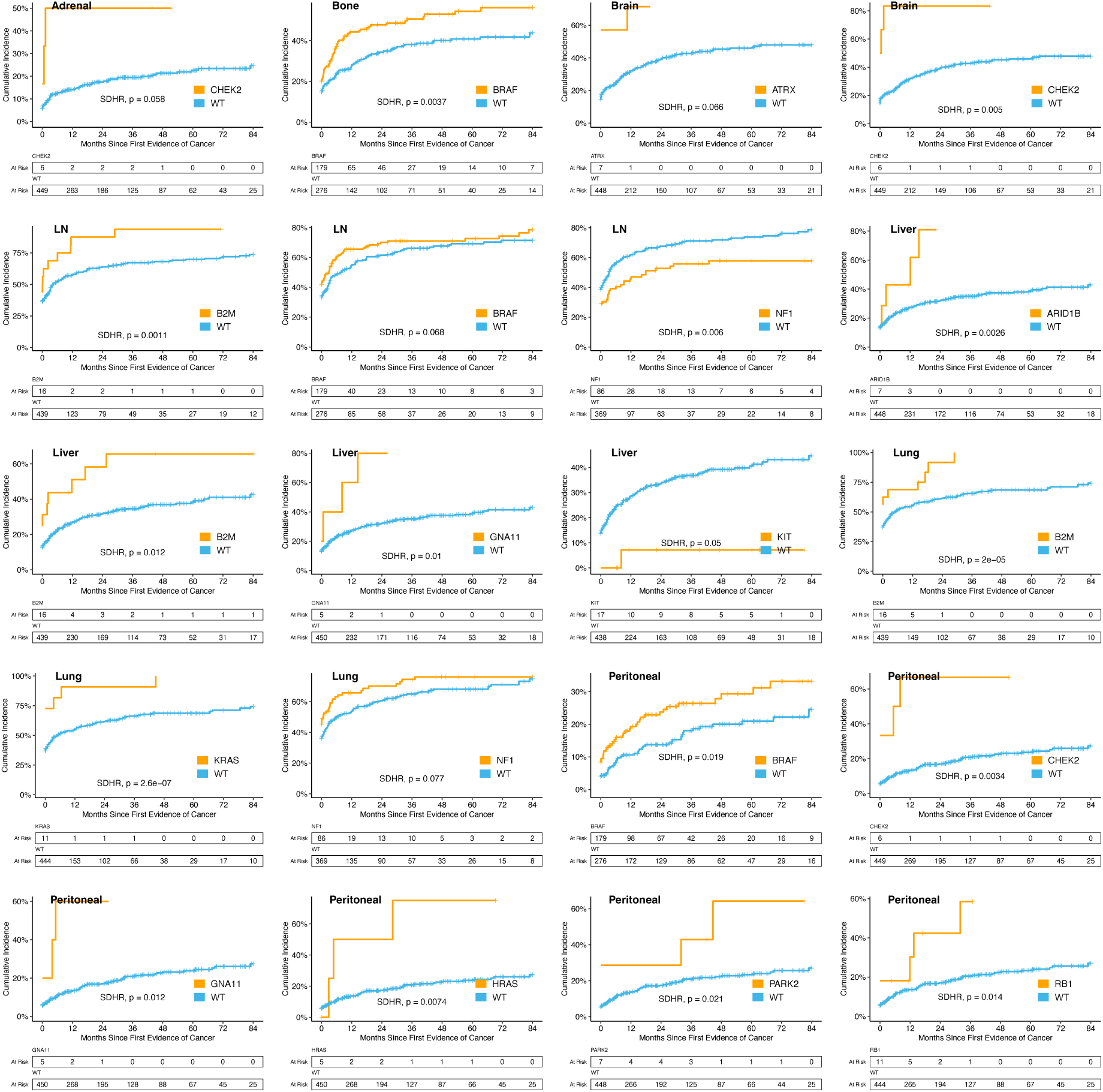
Additional significant CIFs of site-specific metastasis. Cumulative incidence curves of metastasis for specific sites stratified by functional mutations in specific genes. SDHR = subdistribution hazard ratio, p = nominal p-value of subdistribution beta term for gene mutations.

**Supplemental Figure 9.**
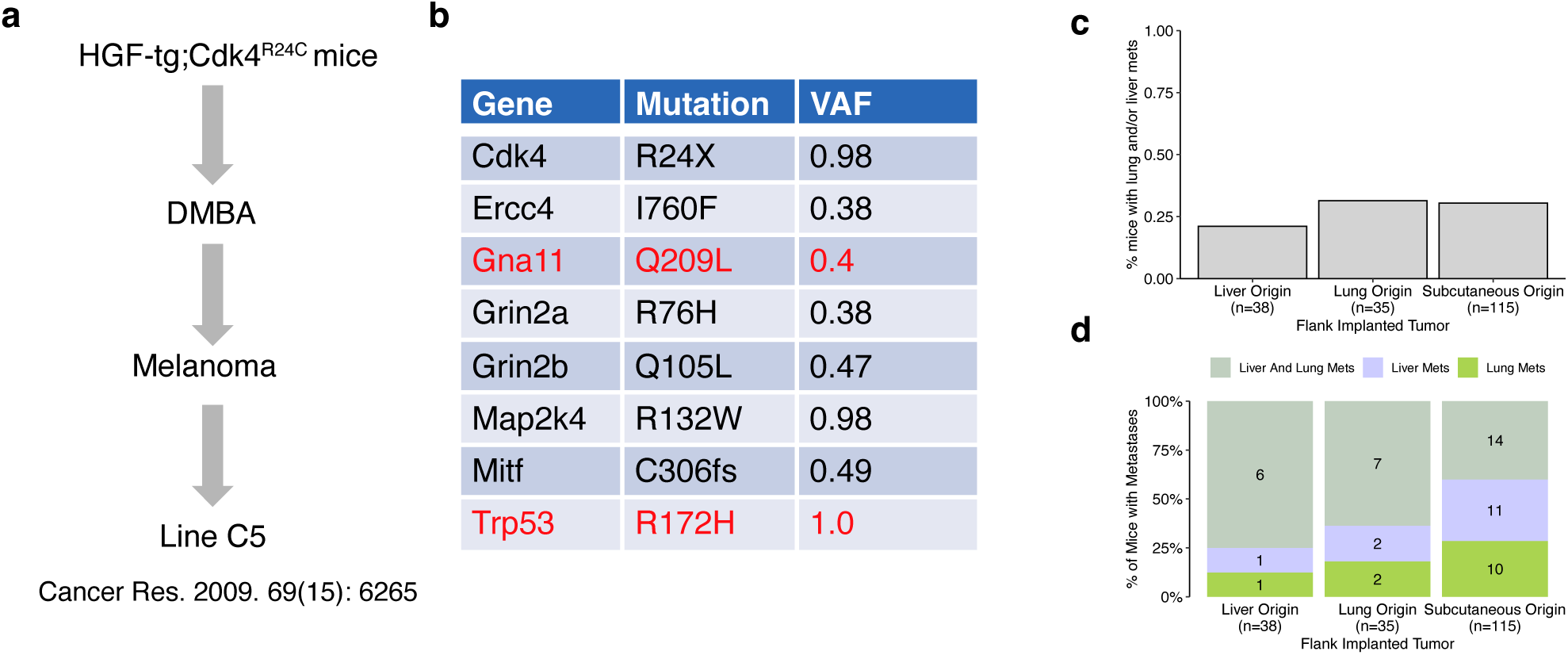
Mouse model generation and further experiment results. **a.** Schematic for the generation of the C5 melanoma cell line. **b.** Results of genomic sequencing of the C5 melanoma cell line. **c,d.** Results from repeating the experiment shown in Figure 7d-e. **c.** Bar plots showing the percentage of mice that developed liver and/or lung metastases after being implanted with tumors originating from different sites. **d.** Stacked bar plots showing the proportions of mice that developed either liver metastases only, lung metastases only, or both lung and liver metastases. Annotation of n refers to the total mice implanted with tumors from the respective origin site.

## Notes

### Summary of Updates

Updated introduction and discussion to make findings and implications more clear. Small modifications to figures to better display major findings. Added and fixed author list.

